# In situ identification of pre-hematopoietic stem cells and their niche

**DOI:** 10.64898/2026.10.05.756028

**Authors:** Xinjian Mao, Ning Zhang, Laura Bennett, Xi He, Allison Scott, Kate Hall, Kaitlyn Petentler, Seth Malloy, Shengping Huang, Amanda Kroesen, Sean McKinney, Athanasios Ploumakis, Zhe Yang, Ruochen Dong, Jose Javier, Jeff Haug, Hua Li, Zulin Yu, Anoja Perera, Ioannis Vlachos, Fei Chen, Paul Trainor, Nancy A. Speck, Linheng Li

## Abstract

Definitive hematopoietic stem cells (HSCs) emerge predominantly along the ventral surface of the dorsal aorta in the embryonic aorta-gonad-mesonephros (AGM) region, yet the spatial organization of the niche underlying this developmental asymmetry remains poorly understood. Defining the native niche of pre-HSCs has also been hindered by their rarity, transient nature and lack of a single definitive marker for their identification *in situ*. Here, we integrated single-cell and spatial transcriptomics to construct a spatiotemporal atlas of the niche in the AGM. Neural cells and chondrogenic mesenchyme predominated dorsally, whereas N-cadherin⁺ mesenchymal stromal cells (N-cad^+^ MSCs) formed a ventral layer between the aortic endothelium and mesonephric cells. BMP and SHH signaling showed overall dorsal enrichment, whereas WNT, NOTCH and KIT signaling were enriched ventrally, with these cellular and signaling patterns dynamically remodeled across HSC ontogeny. Imaging-based spatial transcriptomics validated the findings and further enabled the identification of individual pre-HSCs *in situ* and revealed a multicellular niche comprising endothelial cells, N-cad^+^ MSCs, intra-aortic hematopoietic cluster cells and macrophages. TGFβ and NOTCH signals were primarily supplied locally, whereas major sources of BMP and WNT were more distant. Functional perturbation identified niche-derived CD200 as an immune-regulatory signal promoting HSC development, validating a spatially predicted niche interaction. Together, our findings link tissue-scale spatial organization with the local multicellular niche supporting the preferential development of HSCs along the ventral aorta.

## Introduction

Hematopoietic stem cells (HSCs) sustain lifelong hematopoiesis and form the basis of curative transplantation therapy for numerous hematological disorders^1^. Definitive HSCs first emerge during embryogenesis in the aorta-gonad-mesonephros (AGM) region through a series of conserved developmental transitions^2^. A specialized subset of endothelial cells lining the dorsal aorta first acquires hemogenic identity to generate hemogenic endothelial cells (HECs)^2,3^. HECs subsequently undergo endothelial-to-hematopoietic transition (EHT) to produce intra-aortic hematopoietic clusters (IAHCs), from which only a small subset of cells acquires pre-HSC identity and ultimately matures into transplantable definitive HSCs^4–7^. The intrinsic molecular programs governing these developmental transitions have been extensively investigated^5,8^, leading to the identification of key transcription factors, including RUNX1^6,9^, GATA2^10,11^, HLF^12^, and GFI1^13^, as well as essential developmental signaling pathways, such as NOTCH, BMP, WNT, KIT and inflammatory signaling^11,14–17^. In contrast, how the surrounding niche coordinates these developmental transitions remains largely undefined^18^.

Previous studies have identified several niche cell populations that contribute to embryonic hematopoiesis, including endothelial cells^19^, mesenchymal stromal cells (MSCs)^20–22^, macrophages^23–25^, smooth muscle cells^26^, and neural-derived cells^27,28^. However, how these diverse niche populations are spatially organized to support HSC development remains poorly defined. This question is particularly relevant because EHT occurs primarily on the ventral, rather than the dorsal, side of the dorsal aorta^29–31^. This ventral bias may arise from intrinsic molecular differences between dorsal and ventral endothelial cells, the unique organization of the ventral niche, or both. Early functional studies demonstrated that ventral AGM tissues promote HSC activity, whereas dorsal AGM tissues suppress HSC activity^31^. Recent RNA tomography, microdissection and transcriptomic analyses further revealed dorsoventral molecular asymmetry within the AGM and identified the endothelial-derived regulator EDN1 (Endothelin-1)^19^ and the ventrally enriched regulators BMPER^14^ and ADM^32^. Together, these studies establish functional and molecular asymmetry across the dorsoventral axis of the AGM and identify individual cell populations and signals that contribute to HSC development. However, they do not resolve how these components are assembled into a spatially organized niche, how local niche organization relates to the broader tissue architecture of the AGM, or how these cellular and signaling environments are remodeled across successive stages of HSC development.

A longstanding question in embryonic hematopoiesis is why only a small fraction of HECs undergoing EHT ultimately acquire pre-HSC identity whereas most differentiate into hematopoietic progenitor cells. Although intrinsic heterogeneity of HECs may contribute to this developmental divergence, acquisition of HSC fate may also depend on a pre-HSC-specific niche. Defining this specialized niche requires in situ visualization of individual pre-HSCs. However, pre-HSCs are rare and transient developmental intermediates that lack a single definitive marker and instead are identified by the combinatorial expression of multiple transcription factors and surface markers^6,8,11,12,33–35^. Consequently, they cannot be reliably localized by conventional low-throughput immunofluorescence staining. Here, we sought to define how the spatiotemporal organization of the AGM shapes the specialized niche supporting stage-specific HSC development. We integrated single-cell transcriptomics with complementary sequencing- and imaging-based spatial approaches to resolve the cellular and signaling architecture of the AGM and define the native microenvironment of pre-HSCs in situ.

## Results

### A single-cell atlas of the niche in the AGM

To determine how the niche in the AGM coordinates successive stages of HSC development, we profiled AGM cells across three developmental stages spanning HSC ontogeny. Although hemogenic specification, EHT, and definitive HSC maturation initiate sequentially at approximately embryonic day (E) 9.5, E10.5 and E11.5, the corresponding hematopoietic intermediates, HECs, pre-HSCs and HSCs, reach their peak abundance approximately one day later. We therefore performed single-cell RNA sequencing (scRNA-seq) of all viable cells isolated from E10.5, E11.5 and E12.5 mouse AGM and trunk tissues enabling unbiased profiling of the AGM microenvironment. An E12.5 fetal liver reference sample was also included to facilitate annotation of definitive HSC populations. Because HECs, IAHCs and pre-HSCs are too rare to be comprehensively captured by unenriched sampling, we integrated our dataset with a published scRNA-seq dataset enriched for these populations^36^, generating a unified transcriptional atlas spanning both candidate niche populations and hematopoietic intermediates. Unsupervised clustering identified 38 transcriptionally distinct cell clusters together with 6 hematopoietic developmental populations from the reference dataset (Fig. 1a and Extended Data Fig. 1a). Based on established marker genes, these populations were grouped into four major cellular compartments: endothelial populations expressed canonical vascular markers including *Cdh5* and *Pecam1*; mesenchymal populations were characterized by *Pdgfra* and *Pdgfrb*; neural populations expressed *Ngfr, Sox10*, and *Gata3*; whereas hematopoietic populations expressed *Ptprc, Gypa* and other lineage-specific hematopoietic markers (e.g., *Pf4, Spi1, Hba*) (Fig. 1b and Extended Data Fig. 1b). The relative abundance of individual cell populations changed markedly across the three developmental stages (Extended Data Fig. 1c). Cellular diversity also increased from E10.5 to E12.5, with additional clusters detected at later stages that were not present at E10.5 (Extended Data Fig. 1c), consistent with progressive differentiation and diversification in the AGM throughout HSC ontogeny. Together, these analyses establish a comprehensive single-cell transcriptional atlas encompassing both candidate niche populations and rare hematopoietic intermediates in the AGM throughout HSC development.

**Fig. 1.**
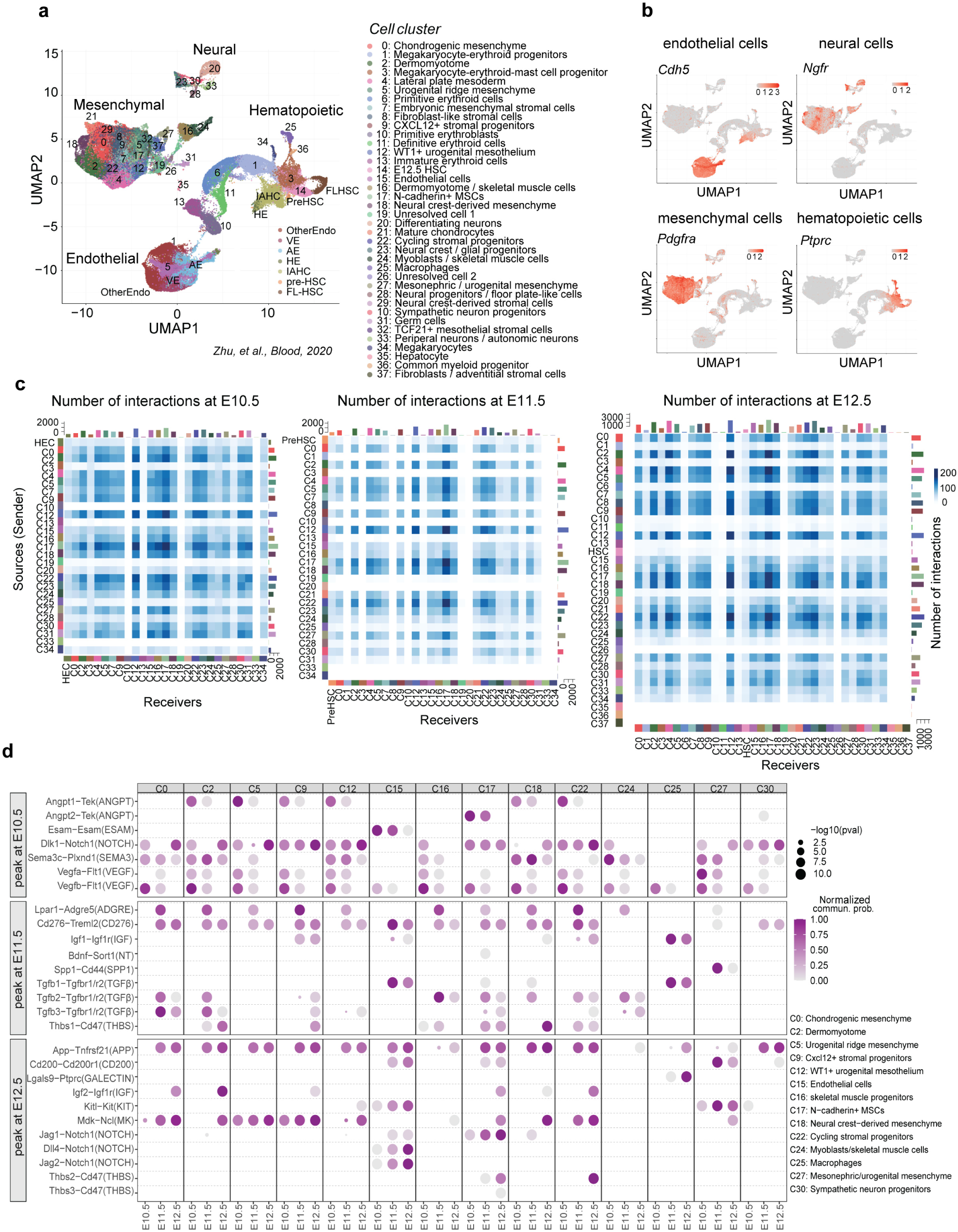
A single-cell atlas of the niche in the AGM. **a,** Uniform Manifold Approximation and Projection (UMAP) visualization of integrated single-cell transcriptomes comprising the published hematopoietic reference dataset enriched for HECs, IAHCs, pre-HSCs and E14.5 fetal liver HSCs (FL-HSCs), together with newly generated datasets from E10.5, E11.5 and E12.5 trunk; E10.5, E11.5 and E12.5 AGM; and E12.5 fetal liver. All viable cells were profiled from the trunk and AGM samples. The E12.5 fetal liver samples comprised one-third CD45⁺LSK (lineage⁻SCA1⁺c-KIT⁺) hematopoietic stem and progenitor cells and two-thirds Ter119⁻ cells. n = 3-5 biologically independent embryonic samples for each newly generated group. **b,** Feature plots showing representative marker genes defining the four major cellular compartments, including endothelial (*Cdh5*), neural (*Ngfr*), mesenchymal (*Pdgfra*) and hematopoietic (*Ptprc*) cells. **c,** Cell-cell communication networks inferred from ligand-receptor interactions at E10.5, E11.5 and E12.5. Heatmaps show the number of predicted interactions between sender and receiver cell populations at each developmental stage. **d,** Representative ligand-receptor interactions enriched at different developmental stages. Dot plots show the predicted communication probability (color) and statistical significance (dot size) of interactions between selected niche populations and hematopoietic intermediates. Receiver cells are HECs at E10.5, pre-HSCs at E11.5, and HSCs at E12.5. Signaling pathways are grouped according to the developmental stage at which they exhibit peak activity. The statistical significance of ligand-receptor interactions was determined using CellChat permutation testing.

We next investigated how niche signaling is dynamically remodeled during HSC development. CellChat analysis identified stage-specific ligand-receptor communication networks and revealed extensive remodeling of cell-cell communication throughout HSC ontogeny. Among all niche populations, Wt1⁺ urogenital mesothelium (cluster 12), N-cadherin⁺ mesenchymal stromal cells (cluster 17; hereafter, N-cad⁺ MSCs) and neural crest-derived mesenchyme (cluster 18) exhibited the strongest interaction activity across all three developmental stages, as measured by both the number and strength of predicted ligand-receptor interactions (Fig. 1c and Extended Data Fig. 1d-f). We next focused on interactions between candidate niche populations and HECs at E10.5, pre-HSCs at E11.5 and HSCs at E12.5 (Fig. 1d). At E10.5, the signaling network displayed a relatively simple organization, with most interactions contributed by several mesenchymal populations, suggesting limited niche specialization. Signaling was enriched for pathways associated with angiogenesis, cell adhesion and early developmental patterning, including *Angpt1, Dlk1, Vegfa* and *Sema3* (a regulator of cell migration and angiogenesis) signaling (Fig. 1d). As development progressed to E11.5 and E12.5, both the number and diversity of signaling pathways increased substantially. Moreover, individual signaling pathways became preferentially associated with specific niche populations rather than being broadly distributed, suggesting increasing specialization of the predicted niche interactions. At E11.5, when the number of pre-HSCs has increased compared to E10.5, and type II (CD45^+^) pre-HSCs are present, IGF and TGFβ signaling became prominent in stromal populations, such as Wt1⁺ urogenital mesothelium, N-cad⁺ MSCs, and Cxcl12⁺ stromal progenitors, together with mesonephric mesenchyme-derived *Spp1* (encoding osteopontin) signaling, and immune-regulatory signals such as *Thbs1* (Thrombospondin1) and *Cd276* (B7-H3) (Fig. 1d). By E12.5, when the AGM contains approximately three definitive HSCs (compared to one at E11.5)^37^, increased KIT, NOTCH, *Mdk* (Midkine, a heparin-binding growth factor involved in cell survival and tissue remodeling), and additional immune-regulatory signaling, including *Cd200* and *Thbs2/3*, accompany a shift toward macrophages, endothelial cells and differentiated stromal populations (Fig. 1d). Together, these analyses establish a stage-resolved ligand-receptor interaction atlas and suggest that dynamic remodeling of the niche in the AGM provides distinct signaling environments for hematopoietic intermediates at successive stages of HSC development.

### Spatiotemporal organization of the AGM tissue and cellular landscape

To define the spatial organization of the HSC niche in the AGM, we performed sequencing-based spatial transcriptomics (Slide-seq V2 and Curio Seeker)^38^ on transverse sections of E10.5, E11.5 and E12.5 mouse embryonic trunks containing the AGM region. Unlike imaging-based spatial transcriptomics approaches that require predefined gene panels, Slide-seq V2 and Curio Seeker enable unbiased genome-wide detection of spatially localized genes. Following quality control, the tissue sections were profiled by densely packed 10-µm barcoded beads that collectively reconstructed genome-wide spatial gene expression at near-cellular resolution. Individual beads detected a median of approximately 500 genes and 1,000 unique molecular identifiers (UMIs) (Extended Data Fig. 2a, b). Unsupervised clustering identified 28 bead clusters, including mesenchymal, endothelial, neural, epithelial and hematopoietic cell clusters (Extended Data Fig. 2c-e). Projection of these clusters back onto tissue sections reconstructed the anatomical organization of the embryonic trunk, including the AGM, neural tube and fetal liver (Extended Data Fig. 2f-h). However, the limited transcript capture per bead reduces cell-type resolution (28 clusters compared with 44 clusters in the scRNA-seq dataset; Fig. 1a), making it difficult to distinguish transcriptionally similar cell populations and accurately resolve the cellular sources of ligands. To overcome this limitation, we projected the scRNA-seq-defined cell clusters in Fig. 1a onto individual beads based on transcriptomic similarity. Most assigned beads were classified as a single cell type, supporting robust spatial annotation of the reconstructed atlas (Extended Data Fig. 3a). This integration leveraged the complementary strengths of scRNA-seq for comprehensive cell-type annotation and Slide-seq V2 / Curio Seeker for spatial information, enabling transcriptionally defined cell populations to be mapped within intact tissue.

The integrated spatial atlas faithfully recapitulated the overall tissue architecture of the embryonic trunk, revealing a highly organized cellular landscape within the AGM (Fig. 2a, b and Extended Data Fig. 3b-l). On the dorsal side of the aorta, chondrogenic mesenchyme formed the dominant stromal compartment immediately adjacent to the aorta, consistent with strong expression of the chondrogenic marker *Pax1* (Fig. 2c, d). More dorsally, neural crest-derived mesenchyme and neural tissues occupied the region beneath and adjacent to the neural tube, in line with expression of canonical neural markers (Extended Data Fig. 3c, d). Endothelial cells were distributed along the aortic wall but did not reconstruct a continuous endothelial monolayer, reflecting the limited resolution of the integrated spatial atlas for fine single-cell-layer structures (Extended Data Fig. 3g, h). As expected, erythroid and other hematopoietic cells were predominantly localized within the aortic lumen (Extended Data Fig. 3i, j). Between the dorsal mesenchyme and the ventral mesonephric compartment, an intermediate stromal region contained abundant *Cxcl12⁺* stromal cells that circumferentially surrounded the dorsal aorta, together with a neural cell population located on both sides of the aorta (Fig. 2e-h). In contrast, the ventral aspect of the AGM displayed substantially greater cellular complexity, including *Wt1⁺* urogenital mesothelium, macrophages, and germ cells (Fig. 2i-n and Extended Data Fig. 3k, l). In addition, we identified a mesenchymal population forming a discrete stromal layer immediately beneath the ventral endothelial lining while being largely absent from the dorsal side, which we refer to here as N-cad⁺ MSCs (Fig. 2m). Notably, this N-cad⁺ MSC layer was positioned between the aortic endothelium and the underlying mesonephric compartment, revealing a layered cellular organization along the ventral aorta. Thus, the ventral aorta is embedded within a cellular environment that is spatially organized into distinct peri-aortic and mesonephric compartments.

**Fig. 2.**
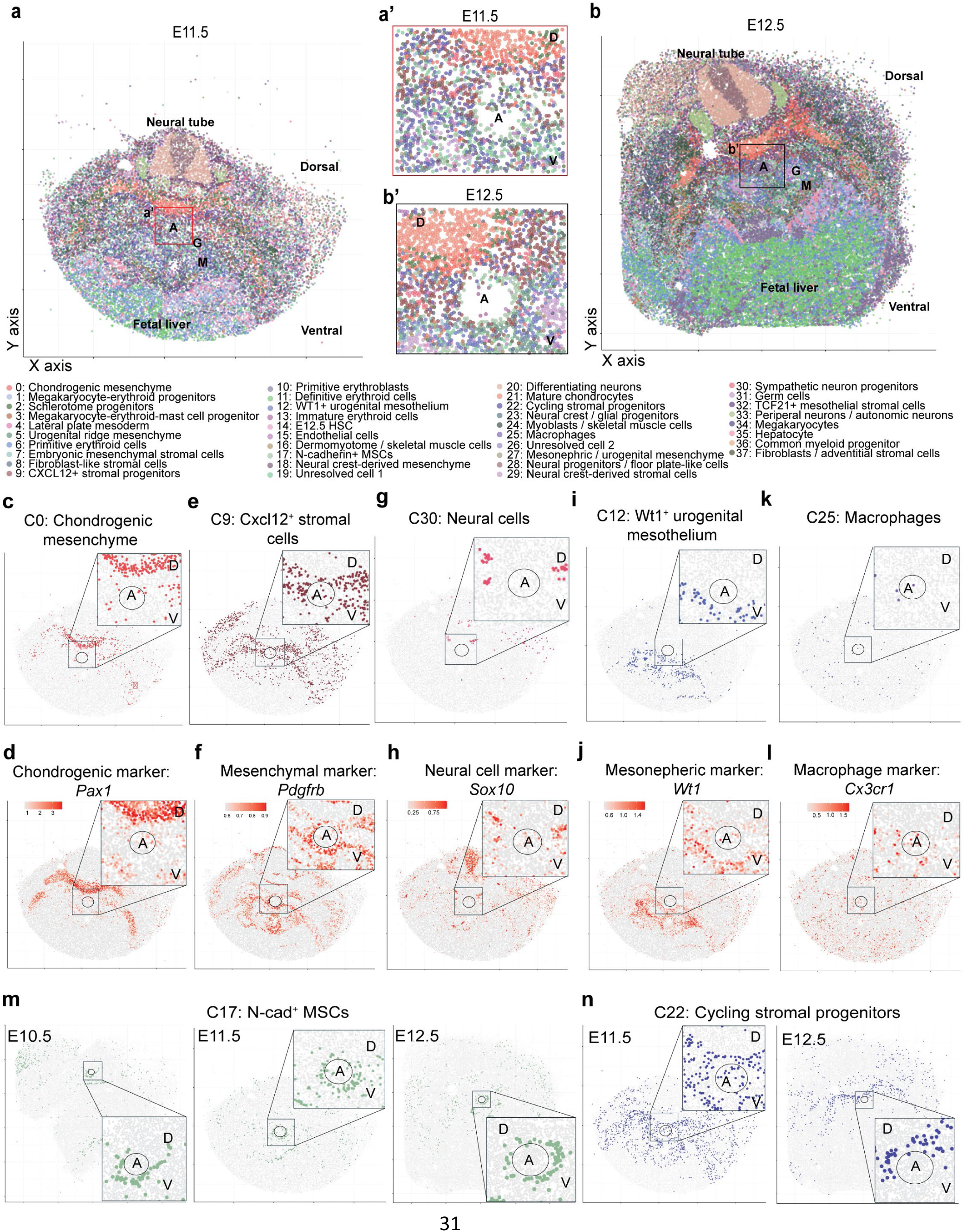
Spatial reconstruction reveals the dynamic organization of the niche in the AGM. **a, b,** Integrated spatial atlas of the embryonic trunk at E11.5 (**a**) and E12.5 (**b**) generated by projecting transcriptionally defined cell populations onto Slide-seq V2 (a) and Curio Seeker (b) sections, as described in Methods. Colors indicate the annotated cell clusters shown in Fig. 1a. Insets (**a′** and **b′**) show enlarged views of the AGM region surrounding the dorsal aorta. For illustrative purposes, we removed erythroid cells in (**a′** and **b′**) to better visualize the aortic structure. n = 3 pucks for the E10.5 and E12.5 groups, and n = 2 pucks for the E11.5 group. Representative sections are shown for each developmental stage. **c-l,** Spatial localization of representative cell populations and their corresponding marker genes within embryonic trunk sections at E11.5, with enlarged view of the AGM region surrounding the dorsal aorta. Shown are chondrogenic mesenchyme (C0; **c**) and *Pax1* expression (**d**), *Cxcl12*⁺ stromal cells (C9; **e**) and *Pdgfrb* expression (**f**), neural cells (C30; **g**) and *Sox10* expression (**h**), *WT1*⁺ urogenital mesothelium (C12; **i**) and *Wt1* expression (**j**), and macrophages (C25; **k**) and *Cx3cr1* expression (**l**). **m,** Spatial distribution of N-cad⁺ MSCs (C17) within embryonic trunk sections at E10.5, E11.5 and E12.5, showing persistent ventrally enriched localization beneath the dorsal aorta throughout HSC ontogeny. **n,** Spatial distribution of cycling stromal progenitors (C22) within embryonic trunk sections at E11.5 and E12.5, illustrating dynamic spatial remodeling during AGM development. A, aorta; G, gonad; M, mesonephros; D, dorsal; V, ventral. The circle indicates the estimated position of the dorsal aorta.

The integrated spatial atlas also enabled temporal characterization of cell dynamics throughout HSC development. For example, N-cad⁺ MSCs persisted as a ventrally enriched stromal layer from E10.5 to E12.5 (Fig. 2m). By contrast, cycling stromal progenitors were absent at E10.5, appeared around the entire aorta at E11.5, and became largely restricted to the dorsal side by E12.5 (Fig. 2n). Together, these analyses establish a spatiotemporal atlas of the niche in the AGM, defining both its cellular architecture and developmental remodeling. However, owing to their rarity, hematopoietic intermediates, including HECs, pre-HSCs and definitive HSCs, could not be confidently resolved by the integrated spatial atlas.

### Spatiotemporal organization of developmental signaling in the AGM

The integrated spatial atlas provides a framework for characterizing both the spatial organization and temporal remodeling of developmental signaling during HSC ontogeny. We first used the E11.5 dataset, corresponding to the peak of EHT and pre-HSC generation, to illustrate the spatial organization of niche signaling in the AGM. It revealed that developmental signaling is highly polarized around the dorsal aorta. For example, BMP signaling exhibited pronounced dorsoventral compartmentalization (Fig. 3a). Whereas *Bmp2, Bmp7* and *Bmper* were predominantly detected in neural cells in dorsal tissues, *Bmp4* was enriched in ventral mesenchymal and mesonephric compartments (Fig. 3b-d and Extended Data Fig. 4e). This indicates that distinct BMP ligands collectively establish a spatially heterogeneous signaling landscape and originate from cells not immediately adjacent to the dorsal aortic wall. In contrast, although a complex WNT signaling pathway network was predicted between pre-HSCs and multiple cell populations around the aorta, visualization of individual WNT ligands revealed that the WNT ligands were predominantly localized to the ventral mesonephric compartment (Extended Data Fig. 4a-d). Thus, WNT and BMP signaling ligands showed broadly opposing distributions across the dorsoventral axis of the dorsal aorta, a classical feature of developmental tissue patterning.

**Fig. 3.**
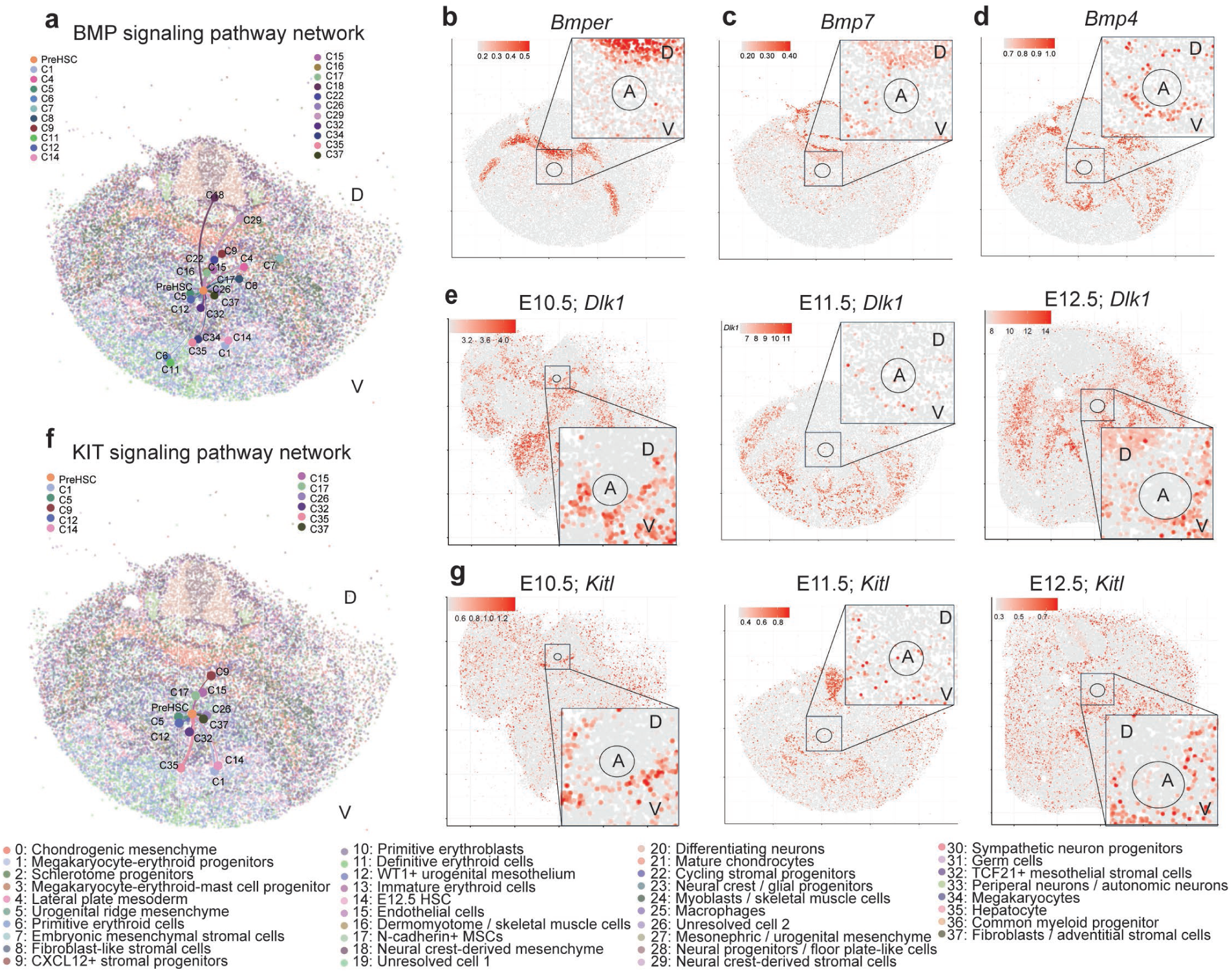
The spatiotemporal organization of developmental signaling in the AGM. **a, f,** The integrated spatial atlas predicted spatially resolved interactions between cell populations and pre-HSCs via the BMP signaling network (**a**) and the KIT signaling network (**f**) within embryonic trunk sections at E11.5. Colored dots indicate ligand-producing cell populations. The position of each dot represents the center of the spatial distribution of the corresponding cell population in the Slide-seq V2 section. Line thickness indicates the predicted interaction strength. **b-d,** Spatial distribution of representative BMP ligands within embryonic trunk sections at E11.5, including *Bmper* (**b**), *Bmp7* (**c**) and *Bmp4* (**d**). **e,** Spatial distribution of *Dlk1* within embryonic trunk sections at E10.5, E11.5 and E12.5. **g,** Spatial distribution of *Kitl* within embryonic trunk sections at E10.5, E11.5 and E12.5. A, aorta; G, gonad; M, mesonephros; D, dorsal; V, ventral. The circle indicates the estimated position of the dorsal aorta.

Beyond these canonical developmental pathways, we also identified numerous signaling molecules with highly restricted anatomical distributions. For example, consistent with previous studies^19^, *Edn1,* an endothelial-derived regulator of embryonic HSC development, was broadly distributed throughout the aortic endothelium (Extended Data Fig. 4g). By contrast, the UGRP1 (Uteroglobin-related protein 1) signaling pathway, an immune-regulatory signaling axis, was enriched within the ventral niche (Extended Data Fig. 4h). *Cxcl12,* a key chemokine that supports HSC maintenance and localization^39^, and *Mdk*, were predominantly expressed by *Cxcl12⁺* stromal cells surrounding the dorsal aorta (Extended Data Fig. 4i, j). *Vip* (vasoactive intestinal peptide), a neuropeptide that has recently emerged as a neuroimmune regulatory molecule in the brain-gut axis^40^, was likewise enriched in the periaortic niche and was predominantly expressed by neural cells (Extended Data Fig. 4k). Together, these findings expand the repertoire of candidate niche-derived factors potentially contributing to embryonic HSC development.

Niche-derived signaling was dynamically remodeled across developmental stages. For example, *Dlk1*, an inhibitory ligand of NOTCH signaling, and *Jag1*, a canonical ligand that activates NOTCH signaling^41^, were primarily derived from endothelial cells and N-cad⁺ MSCs progressively increased from E10.5 to E12.5 (Fig. 3e and Extended Data Fig. 5a). *Kitl* expression was likewise enriched within the ventral aorta (Fig. 3f, g) and progressively increased from E10.5 to E12.5, consistent with its established role in adult HSC expansion and maintenance^42^. In addition, several niche-derived factors associated with neurotrophic signaling (*Bdnf,* brain-derived neurotrophic factor), extracellular matrix remodeling (*Thbs2*), and stem cell niche regulation (*Spp1*) also progressively increased from E10.5 to E12.5 (Extended Data Fig. 5b-d). Together, these analyses demonstrate that the signaling environment surrounding the dorsal aorta is organized across both anatomical space and developmental time, with regional developmental signals and peri-aortic niche signals displaying distinct spatial distributions throughout HSC ontogeny.

### Single-cell spatial imaging validates the niche in the AGM

Unlike the integrated spatial atlas, which relies on computational reconstruction, imaging-based spatial transcriptomics directly measures endogenous transcripts at single-cell resolution, providing an approach to validate and refine the integrated spatial atlas. We therefore performed multiplexed error-robust fluorescence in situ hybridization (MERFISH)^43^ on E11.5 embryonic trunk sections using a customized panel of 140 genes comprising cell markers together with selected ligands and receptors (Extended Data Table 1). The MERFISH data faithfully reconstructed the overall anatomy of the embryonic trunk, with particularly detailed reconstruction of the AGM region owing to the enrichment of the probe panel for genes marking niche cells and signals surrounding the dorsal aorta (Fig. 4a and Extended Data Fig. 6a, b). It confirmed the polarized cellular organization while resolving anatomical features that could not be faithfully reconstructed by the integrated spatial atlas. For example, endothelial cells formed a continuous luminal monolayer lining the dorsal aorta (Fig. 4b), consistent with previous immunofluorescence studies^19,22^ and providing substantially improved single-cell spatial resolution compared with that of the integrated spatial atlas (Extended Data Fig. 3g). Immediately beneath the ventral, but not dorsal, endothelium, N-cad⁺ MSCs formed a discrete second stromal layer, confirming their ventrally enriched distribution. Importantly, MERFISH further resolved N-cad⁺ MSCs between the aortic endothelial lining and the adjacent mesonephric cells, independently confirming the layered cellular organization. Consistent with this data, the area dorsal to the dorsal aorta was primarily occupied by notochord-, chondrogenic- and neural-associated tissues, whereas the ventral side contained a considerably more complex mesenchymal and mesonephric compartment (Fig. 4a, b). Together, these findings independently validate the tissue and cellular organization in the AGM while refining anatomical features that could not be fully resolved by the integrated spatial atlas.

**Fig. 4.**
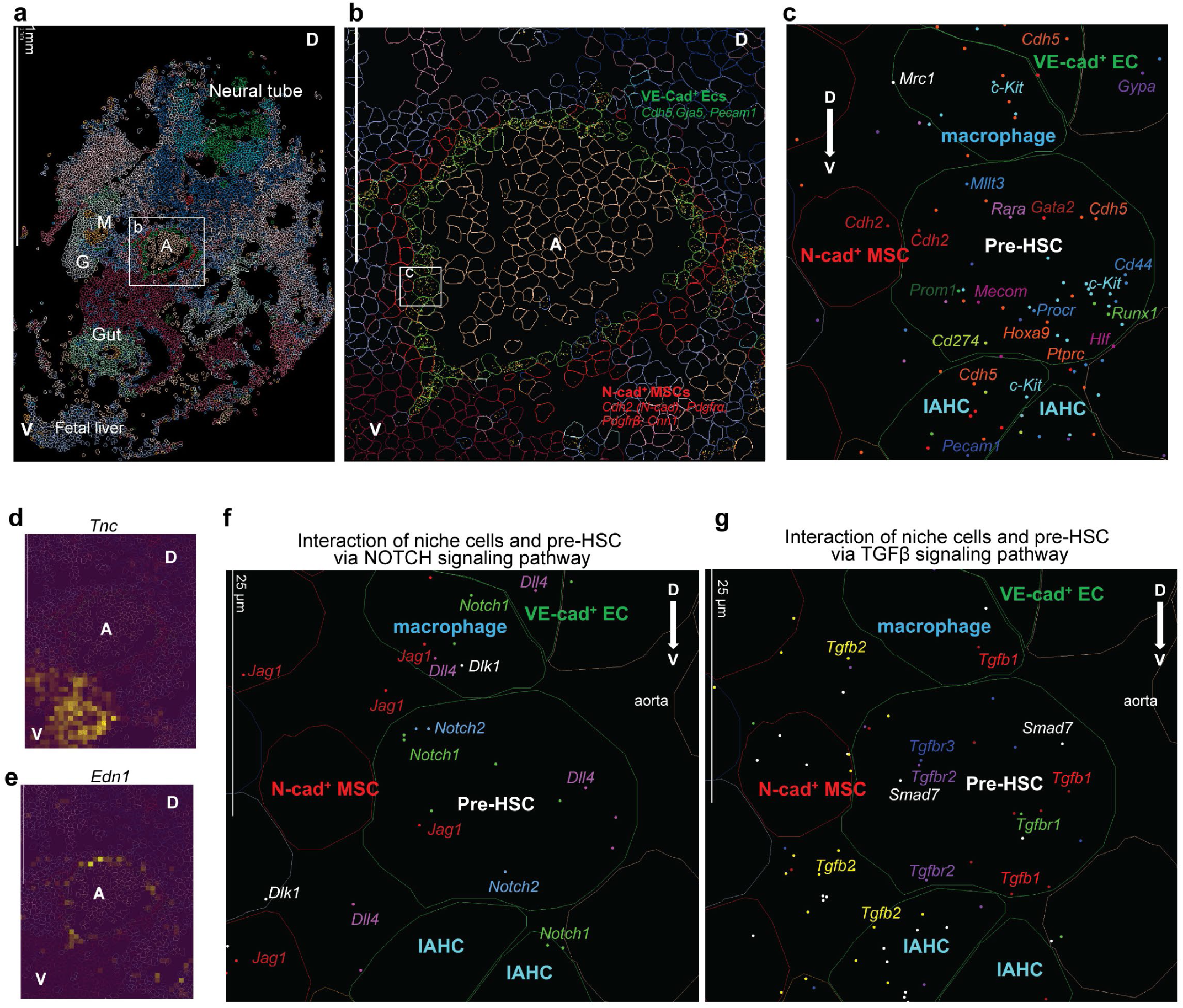
In situ identification of pre-HSCs and their native multicellular niche. **a,** MERFISH reconstruction of an E11.5 embryonic trunk section. The boxed region indicates the dorsal aorta shown at higher magnification in **b**. Representative images are shown from n = 18 biologically independent embryonic trunk sections. **b,** High-magnification view of the AGM showing the cellular organization surrounding the dorsal aorta. Aortic endothelial cells (VE-cad⁺ ECs, expressing *Cdh5, Gja5*, and *Pecam1*) form a continuous luminal monolayer, whereas N-cad⁺ MSCs (expressing *Cdh2, Pdgfra, Pdgfrb*, and *Cnn1*) form a discrete stromal layer predominantly beneath the ventral endothelium. Each dot represents an individual EC-specific transcript (*Cdh5, Gja5*, or *Pecam1*) detected in situ, demonstrating the specificity of transcript detection and the robustness of EC identification. The boxed region indicates the pre-HSC and its surrounding cells shown at higher magnification in **c**. **c,** Representative pre-HSC identified in situ by multiplex detection of pre-HSC-enriched transcripts. The pre-HSC co-expresses characteristic transcription factors and surface markers, including *Runx1, Gata2, Mecom, Mllt3, Hlf, Procr, Kit, Cd44, Cd274, Prom1, Hoxa9* and *Ptprc*, and is positioned adjacent to aortic endothelial cells, N-cad⁺ MSCs, IAHC cells (expressing *Cdh5* and *c-Kit*) and an *Mrc1^+^* macrophage. **d, e,** Representative spatial expression of candidate niche signal *Tnc* (**d**) and *Edn1* (**e**) within the embryonic trunk section. **f,** Representative MERFISH image illustrating ligand-receptor interactions between neighboring niche cells and a pre-HSC through the NOTCH signaling pathway. Ligands (*Jag1, Dll4* and *Dlk1*) and receptors (*Notch1* and *Notch2*) are shown in their endogenous spatial locations. **g,** Representative MERFISH image illustrating ligand-receptor interactions between neighboring niche cells and a pre-HSC through the TGFβ signaling pathway. Ligands (*Tgfb1-3*), receptors (*Tgfbr1-3*) and the downstream target *Smad7* are shown in their endogenous spatial locations. A, aorta; G, gonad; M, mesonephros; D, dorsal; V, ventral. ECs, endothelial cells. Each dot in (**b, c, f, g**) represents an individual transcript detected in situ. Scale bar = 1 mm in (**a**) and 25 µm in (**f, g**).

The MERFISH data also validated and refined the spatial organization of developmental signaling. Consistent with the integrated spatial atlas, NOTCH signaling ligands, *Dll4* and *Jag1*, were distributed around the aorta, while *Dlk1* was highly enriched on the ventral side of the aorta (Extended Data Fig. 6c). BMP signaling also exhibited pronounced spatial compartmentalization. *Bmp2* was localized immediately adjacent to the dorsal aortic wall, suggesting production by endothelial or peri-aortic mesenchymal cells, whereas *Bmp7* was restricted to more distant dorsal neural tissues (Extended Data Fig. 6d). By contrast, *Bmp4* and *Bmper* were enriched within ventral mesenchymal and mesonephric compartments (Extended Data Fig. 6d), refining the projection-based reconstruction and bringing the localization of *Bmper* into agreement with previous functional studies^14^. *Edn1* localized to the endothelial lining without an obvious spatial preference (Fig. 4e), whereas *Shh* (Sonic hedgehog) expression remained confined to the dorsal notochord (Extended Data Fig. 6e). In contrast, KIT- and WNT-associated ligands (Extended Data Fig. 6f, g), together with *Tnc* (Tenascin-C, Fig. 4d), which has been reported to mediate embryonic hematopoiesis^44^, preferentially localized to the ventral mesenchymal compartment. MERFISH validated mesonephric cells as a prominent source of WNT ligands, confirming the association of WNT signaling with the ventral and mesonephric compartments. Together, the sequencing-and imaging-based spatial transcriptomic approaches combined unbiased genome-wide discovery with direct single-cell validation, establishing a robust single-cell spatiotemporal atlas for investigating the ventral bias of definitive HSC emergence.

### In situ identification of pre-HSCs reveals their native multicellular niche

A major unresolved challenge in the field is to define the native microenvironment specifically associated with pre-HSCs, as opposed to other hematopoietic progenitor cells within the IAHCs. This has been difficult because pre-HSCs are rare and transient and lack a single definitive marker for their identification in situ. We therefore leveraged the high-throughput and spatially resolved transcript detection of MERFISH to overcome this limitation. Notably, MERFISH robustly identified individual pre-HSCs in situ for the first time by directly and simultaneously detecting a panel of pre-HSC-enriched transcripts within the same cell, including *Runx1, Gata2, Mecom, Mllt3, Hlf, Procr, Kit, Cd44, Cd274, Prom1, Hoxa9* and *Ptprc*^6,8,11,12,33–35^ (Fig. 4c and Extended Data Fig. 7a, b). Localization of individual pre-HSCs subsequently enabled delineation of their immediate native microenvironment. Rather than being associated with a single supporting cell type, individual pre-HSCs were consistently embedded within a highly organized multicellular niche unit comprising adjacent aortic endothelial cells, N-cad⁺ MSCs, neighboring IAHCs and, in some cases, macrophages (Fig. 4c and Extended Data Fig. 7a, b). Notably, endothelial cells and N-cad⁺ MSCs were also observed along the relatively dorsal side of the aorta, where pre-HSCs and IAHCs were less frequently detected. Therefore, the presence of these two cell populations alone may not fully account for the preferential emergence of HSCs along the ventral aorta. Instead, their organization with additional niche cells, together with broader tissue-level patterning, may contribute to establishing spatially favorable environments for HSC specification.

### The signaling landscape surrounding pre-HSCs

We next dissected the signaling interactions that coordinate HSC development by examining the spatial distributions of ligands and receptors between pre-HSCs and neighboring niche cells. The MERFISH data revealed the spatial organization of multiple signaling components within individual developmental pathways. Rather than being uniformly produced, distinct ligands within the same signaling pathway originated from different niche cell populations, revealing differences in the cellular sources of candidate signals within the multicellular niche. In the NOTCH pathway, *Notch1* and *Notch2* receptors were detected in pre-HSCs, whereas the three major ligands exhibited strikingly distinct cellular origins (Fig. 4f and Extended Data Fig. 7c, d). *Dll4* expression was largely restricted to IAHCs and, notably, pre-HSCs, suggesting that developing HSCs are not merely passive recipients of niche-derived signals but may also actively participate in shaping their local signaling environment^15^. *Jag1* was broadly detected in adjacent N-cad⁺ MSCs, aortic endothelial cells and IAHCs, whereas *Dlk1*, an inhibitory ligand of NOTCH signaling, was predominantly expressed by N-cad⁺ MSCs. Similarly, in the TGFβ pathway, *Tgfbr1-3* and the downstream target *Smad7*, were detected within pre-HSCs (Fig. 4g and Extended Data Fig. 7e, f). *Tgfb1* was largely restricted to IAHCs and pre-HSCs, whereas *Tgfb2* was preferentially enriched in adjacent N-cad⁺ MSCs. Although detected at lower transcript abundance, *Tgfb3* expression was also observed selectively in N-cad⁺ MSCs (Extended Data Fig. 7f), further supporting N-cad⁺ MSCs as a prominent local source of niche signaling. In contrast, components of the BMP and WNT pathways were less frequently detected within the immediate multicellular niche of pre-HSCs (Extended Data Fig. 7g-i). Together with the broader spatial analyses identifying mesonephric and other surrounding tissues as prominent sources of developmental signals, these findings reveal two spatial levels of signaling organization: signals supplied within the immediate multicellular niche (e.g., NOTCH and TGFβ) and signals arising from more distant tissue compartments (e.g., BMP and WNT).

### N-cad⁺ MSCs define a distinct peri-aortic mesenchymal population

Our spatial analyses highlighted N-cad⁺ MSCs as a prominent component of the developing HSC niche. N-cad⁺ MSCs formed a ventrally enriched mesenchymal population and occupied an intermediate position between the aortic endothelium and mesonephric cells, forming a layered cellular organization along the ventral aorta where EHT preferentially occurs (Fig. 2m and Fig. 4b). N-cad⁺ MSCs were predicted to interact extensively with developing hematopoietic populations (Fig. 1d). MERFISH further identified N-cad⁺ MSCs as a component of the multicellular niche surrounding pre-HSCs and revealed their direct contact with pre-HSCs (Fig. 4c and Extended Data Fig. 7a, b). These findings prompted us to characterize N-cad⁺ MSCs in greater detail.

We therefore compared N-cadherin expression with representative mesenchymal markers using both MERFISH data and immunofluorescence staining. Consistent with previous studies^20,22^, *Pdgfra* and *Pdgfrb* broadly labeled mesenchymal populations throughout the AGM, whereas *Cnn1* (encoding Calponin-1) marked vascular smooth muscle cells distributed relatively uniformly around the dorsal aorta (Extended Data Fig. 8a, b). By contrast, *Cdh2* (encoding N-cadherin) marked neural cell populations and a discrete mesenchymal population enriched beneath the ventral aortic endothelium (Extended Data Fig. 8a, b). Although N-cadherin partially overlapped with Calponin-1, *Pdgfra* and *Pdgfrb*, its distinct spatial distribution distinguished N-cad⁺ MSCs from vascular smooth muscle cells and broader peri-aortic mesenchymal populations. N-cadherin therefore provided a more specific marker for this ventrally enriched MSC population.

### Niche-derived CD200 signaling promotes embryonic HSC development

We next sought to determine whether the spatially predicted interactions between niche cells and developing HSCs have functional significance. Our analysis identified immune-regulatory signaling as a previously unrecognized niche component in the AGM. Among these signals, the CD200-CD200R axis provided a candidate for functional validation because CD200 is a membrane-bound ligand that mediates contact-dependent signaling. CellChat identified both N-cad⁺ MSCs and endothelial cells as major sources of CD200, with predicted signaling activity progressively increasing from E10.5 to E12.5 (Fig. 1d). Immunofluorescence confirmed CD200 expression throughout the peri-aortic region surrounding the dorsal aorta (Extended Data Fig. 8c). We therefore selected CD200-CD200R signaling to functionally test a spatially predicted niche interaction.

To determine the functional contribution of niche-derived CD200, we generated two conditional knockout models that targeted *Cd200* deletion to N-cadherin-expressing cells (*N-cadherin-CreER; Cd200 ^fl/fl^*) or endothelial cells (*VE-cadherin-Cre; Cd200 ^fl/fl^*) (Fig. 5a, g), respectively. Efficient CD200 deletion in the targeted populations was confirmed by flow cytometry (Extended Data Fig. 9a-c). Deletion of *Cd200* from either N-cad⁺ MSCs or endothelial cells produced remarkably similar developmental phenotypes. In both models, there were minimal changes in IAHC formation at E10.5 and E11.5 (Fig. 5b-d and Fig. 5h-j). By contrast, transplantation assays consistently demonstrated impaired functional HSC activity, with no significant defects following transplantation of E11.5 AGM cells but substantially reduced multilineage engraftment from E12.5 AGM cells (Fig. 5e, f, k, l, and Extended Data Fig. 9e-h). These stage-dependent functional defects closely paralleled the developmental increase in predicted CD200-CD200R signaling.

**Fig. 5.**
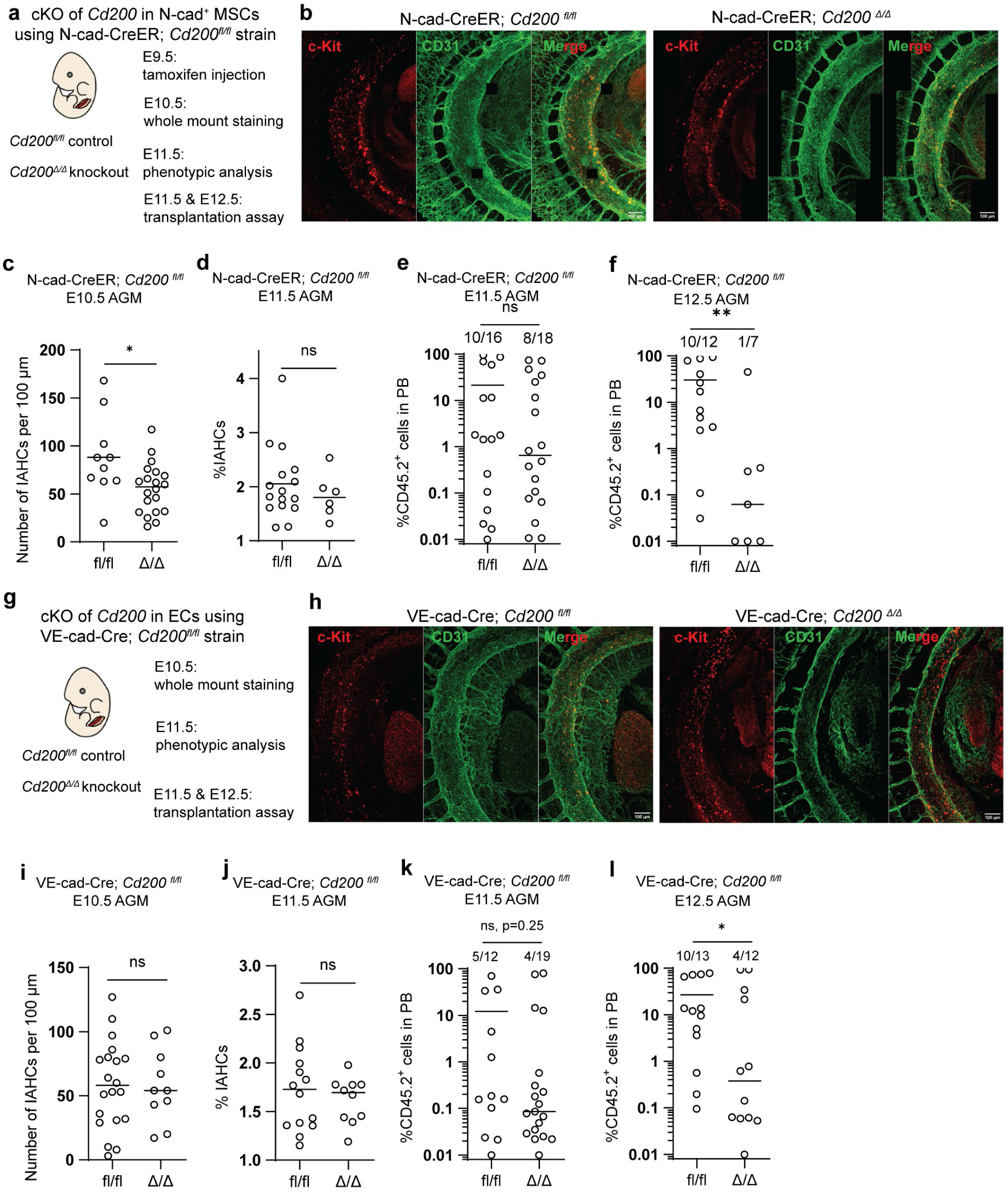
Niche-derived CD200 promotes embryonic HSC development. **a,** Experimental design for conditional deletion of *Cd200* in N-cad⁺ MSCs. *N-cad*-CreER; *Cd200* ^fl/fl^ embryos received tamoxifen at E9.5. Whole-mount immunofluorescence was performed at E10.5, phenotypic analyses of IAHCs at E11.5, and transplantation assays using E11.5 and E12.5 AGM cells **b, c,** Representative whole-mount immunofluorescence images (**b**) and quantification (**c**) of E10.5 embryos stained for c-Kit (red) and CD31 (green) following deletion of *Cd200* in N-cad⁺ MSCs. IAHC cells were quantified as c-Kit^+^ CD31^+^ cells. Representative images are shown from n = 3 biologically independent embryos. **d,** Quantification of phenotypic c-Kit^+^CD31^+^ IAHC cells at E11.5 following deletion of *Cd200* in N-cad⁺ MSCs, as assessed by flow cytometry. n = 16 embryos for fl/fl group and n = 6 embryos for Δ/Δ group across 3 independent pools. **e, f,** Peripheral blood donor chimerism in recipient mice 16 weeks after transplantation of E11.5 (**e,** 1 e.e. per recipient) or E12.5 (**f,** 0.5 e.e. per recipient) AGM cells isolated from littermate control (*Cd200* ^fl/fl^) and *N-cad-*CreER; *Cd200* ^Δ/Δ^ embryos. Each dot represents one recipient mouse. In **e**, n = 16 for fl/fl group and n = 18 for Δ/Δ group across 3 independent pools; In **f**, n = 12 for fl/fl group and n = 7 for Δ/Δ group across 3 independent pools. Fractions indicate the number of mice with successful engraftment, defined as ≥1% donor chimerism. **g,** Experimental design for endothelial-specific deletion of *Cd200* using *VE-cad*-Cre; *Cd200* ^fl/fl^ mice. Whole-mount immunofluorescence was performed at E10.5, phenotypic analyses at E11.5, and transplantation assays using E11.5 and E12.5 AGM cells. **h, i,** Representative whole-mount immunofluorescence images (**h**) and quantification (**i**) of E10.5 embryos stained for c-Kit (red) and CD31 (green) following deletion of *Cd200* in VE-cad⁺ endothelial cells. IAHC cells were quantified as c-Kit^+^ CD31^+^ cells. Representative images are shown from n = 3 biologically independent embryos. **j,** Quantification of phenotypic c-Kit^+^ CD31^+^ IAHCs at E11.5 following endothelial-specific deletion of *Cd200*. Each dot represents one embryo. n = 16 for fl/fl group and n = 6 for Δ/Δ group across 3 independent pools. **k, l,** Peripheral blood donor chimerism in recipient mice 16 weeks after transplantation of E11.5 (**k**, 1 e.e. per recipient) or E12.5 (**l**, 0.5 e.e. per recipient) AGM cells from littermate control (*Cd200* ^fl/fl^) and *VE-cad*-Cre; *Cd200* ^Δ/Δ^ embryos. Each dot represents one recipient mouse. In **k**, n = 12 for fl/fl group and n = 19 for Δ/Δ group across 3 independent pools; In **l**, n = 13 for fl/fl group and n = 12 for Δ/Δ group across 3 independent pools. Fractions indicate the number of mice with successful engraftment, defined as ≥1% donor chimerism. Statistical significance was determined using unpaired t test in (**c, d, i, j**) and Fisher’s exact test in (**e, f, k, l**). e.e., equivalent embryo. cKO, conditional knockout. Lines present means (**c-f, i-l**). *P < 0.05; **P < 0.01; ns, not significant. Scale bar = 100 µm in (**b, h**).

Because CD200-CD200R signaling has been linked to MAPK activation, nitric oxide synthase activation and WNT signaling in a context-dependent manner^45–47^, we next investigated downstream signaling by examining ERK phosphorylation, nitric oxide levels and the nuclear-to-cytoplasmic β-catenin ratio in phenotypic pre-HSCs using the *N-cadherin-CreER; Cd200 ^fl/fl^* model. Conditional deletion of *Cd200* from N-cad⁺ MSCs significantly reduced ERK phosphorylation in pre-HSCs while exerting only a modest effect on nitric oxide levels and the nuclear-to-cytoplasmic β-catenin ratio (Extended Data Fig. 9i-l). These findings suggest that MAPK signaling may represent one downstream pathway mediating the effects of CD200 on HSC development. Together, these two genetic mouse models provide complementary functional validation of the spatially predicted niche interaction and establish CD200 as a physiologically relevant immune-regulatory niche signal that promotes HSC development.

### Establishment of immune-regulatory characteristics during HSC maturation

Adult HSCs possess well-established immune-regulatory and immune-privileged properties. When these characteristics are established during HSC ontogeny remains unclear. Our analyses identified multiple immune-related ligand-receptor interactions that progressively increased during HSC ontogeny (Fig. 1d). We therefore asked whether immune-regulatory characteristics begin to emerge during embryonic HSC development. Analysis of the scRNA-seq dataset revealed progressive upregulation of multiple immune-regulatory genes as cells progressed from HECs through IAHCs and pre-HSCs to HSCs, including *Cd47, Cd274* (encoding PD-L1), *Cd55* (a regulator of C3 and C5 convertases), *Cd59b* (a complement-regulatory protein), *Tap1* and *Tap2* (transporters associated with antigen processing), and *B2m* (β2-microglobulin, an essential structural component of MHC-I complexes) (Extended Data Fig. 10a). To validate these findings at the protein level, we analyzed representative immune-regulatory molecules in type II pre-HSCs from the E12.5 AGM by flow cytometry. Type II pre-HSCs expressed high levels of CD200R1, CD47, PD-L1, CD74 (the invariant chain of MHC-II), and CD73 (an enzyme involved in adenosine production) (Extended Data Fig. 10b-g). These findings indicate that immune-regulatory characteristics begin to emerge during embryonic HSC development.

Because CD200 itself is a well-characterized immune-checkpoint ligand, we next asked whether niche-derived CD200 regulates this broader immune-regulatory program. Conditional deletion of *Cd200* from N-cad⁺ MSCs had little effect on the expression of most immune-regulatory molecules examined, apart from a modest reduction in CD47 expression (Extended Data Fig. 10c-g). These findings suggest that CD200 contributes to HSC development without broadly regulating the acquisition of the immune-regulatory program.

## Discussion

Here we establish a single-cell spatiotemporal atlas of niche cells and their associated signals in the AGM, providing a spatial framework for understanding the long-recognized ventral bias of definitive HSC emergence at single-cell resolution. Previous studies established functional and molecular asymmetry across the dorsoventral axis of the AGM but largely relied on regional microdissection and analyses at individual developmental stages to identify regulators of HSC development^14,19,30–32^. Our study extends this work by resolving the tissue and cellular organization of the AGM beyond a simple dorsoventral division and showing how niche composition and signaling sources are dynamically remodeled across HSC ontogeny (Extended Data Fig. 10h). At the tissue level, we resolved a layered cellular organization along the ventral aorta, with N-cad⁺ MSCs occupying the interface between the aortic endothelium and the mesonephric compartment. This cellular architecture coincided with marked dorsoventral polarization of developmental signals: BMP signaling showed an overall dorsal bias, although several BMP-related factors (e.g., BMP4 and BMPER) were also enriched ventrally, whereas SHH predominated dorsally, and WNT, NOTCH and KIT were enriched ventrally. Notably, the broadly opposing distributions of WNT and BMP resemble classical developmental patterning and analogous signaling gradients in adult stem cell niches, including the intestinal crypt^48,49^. This similarity raises the possibility that opposing WNT-BMP organization represents a recurring spatial principle for establishing stem-cell-supportive niches across developmental and adult tissues.

By combining MERFISH with multiplex detection of pre-HSC-enriched transcripts, we extend previous analyses of heterogeneous IAHCs by identifying individual pre-HSCs in situ and resolving the cellular organization of their immediate native microenvironment. Endothelial cells^19^, macrophages^23–25^ and IAHCs^15,50^ have previously been implicated in embryonic HSC development, and our functional studies establish N-cad⁺ MSCs as an additional niche component, providing functional support for the multicellular organization identified around pre-HSCs. The multicellular nature of this niche was also reflected in its signaling organization, with individual developmental pathways frequently receiving inputs from more than one niche population, consistent with previously reported multicellular sources of inflammatory signaling^50^. Such redundancy may buffer fluctuations in signal availability and provide robustness to the developing HSC niche. Spatial analysis further showed that signaling to pre-HSCs extends beyond their immediate cellular neighbors. For example, NOTCH and TGFβ were predominantly supplied by cells surrounding pre-HSCs, whereas major sources of WNT and BMP resided in more distant tissue compartments. Thus, the signals received by pre-HSCs may reflect integration between their immediate multicellular niche and the more distant tissue environment. These observations raise the possibility that local niche cells provide direct support for HSC development, while broader tissue organization spatially defines where these interactions occur, potentially contributing to the preferential emergence of HSCs within the layered cellular environment formed by aortic endothelial cells, N-cad⁺ MSCs and mesonephric cells.

Our findings also connect immune-regulatory programs to embryonic HSC development. Immune-regulatory molecules were progressively acquired during pre-HSC maturation, consistent with previous studies showing upregulation of immune-response genes (e.g., *Cd244, Cd74, Cd200r4* and *Cd274*) during *ex vivo* maturation of pre-HSCs into functional HSCs^34^. PD-L1 is similarly upregulated during maturation but is dispensable for HSC engraftment in immunodeficient recipients^34^, suggesting that acquisition of these molecules does not necessarily reflect an immediate requirement for immune protection. Our findings identify a functional role for CD200 beyond its association with this broader expression program: its deletion did not substantially alter the immune-regulatory state of pre-HSCs but impaired HSC development, potentially through MAPK signaling. In adult HSCs, PD-L1 has been shown to enhance engraftment after *ex vivo* expansion in allogeneic recipients^51^, whereas CD200 contributes to HSC immune tolerance in bone marrow^47^. These findings raise the possibility that some immune-regulatory molecules acquired during HSC maturation may initially support HSC development and later contribute to immune protection. Determining when and how such developmental and immune-protective functions become linked will be important for understanding the establishment of immune privilege during HSC ontogeny.

Overall, our study extends the concept of the embryonic HSC niche from individual supporting cell types to two interconnected spatial levels: the local multicellular niche surrounding pre-HSCs and the broader tissue organization in which this niche is embedded. This framework may inform efforts to reconstruct the spatial organization of developmental HSC niches *ex vivo* for regenerative applications.

## Acknowledgements

We thank the Broad Institute and Dr. Fei Chen’s group for providing the Slide-seq V2 pucks. We thank the BIDMC Spatial Technologies Unit (STU, RID:SCR_024905) and its personnel for the provision of spatial tissue profiling services. The Spatial Technologies Unit is a BIDMC Institutional Core Facility and a Harvard Cancer Consortium Core. Harvard Cancer Consortium is supported in part by an NCI Cancer Center Support Grant # NIH 5 P30 CA06516. We thank Nikolaos Kalavros from the STU for support in designing the spatial transcriptomics probe set. We thank H. Monnin, E. Hamilton, I. Dunwiddie, S. Billinger, and M. Peterson at Stowers Institute for Medical Research for technical support. We thank Diana Sá da Bandeira at St. Jude Children’s Research Hospital for scientific discussions. We acknowledge the University of Kansas Medical Center’s Genomics Core for sequencing support. P. Trainor was supported by the Stowers Institute for Medical Research (SIMR-1006). L. Li was supported by the Stowers Institute for Medical Research (SIMR-1004). X. Mao was supported by an ASH Scholar Award. N. A. Speck and L. Bennett were supported by 1 R01 HL163265 and R01 HL091724.

## Author contributions

X.M. conceived the project, designed and performed the experiments, analyzed the data, and wrote the manuscript. N.Z. performed the bioinformatic analyses. L.B. provided training in phenotypic and transplantation assays. X.M., A.S., K.H., K.P., and S.M. performed the Slide-seq V2, Curio Seeker, and MERFISH experiments. X.H. provided training and assisted with the experiments. S.H., S.M. and A.K. performed immunofluorescence and whole-mount staining. I.V. and A.P. contributed to the MERFISH assay. F.C. contributed to the Slide-seq V2 assay. Z.Y. and R.D. contributed to animal experiments. J.J. and J.H. contributed to flow cytometry assays. S.M. contributed to MERFISH data analysis. A.P., Z.Y., and H.L. provided technical support. P.T. provided training in AGM dissection and assisted with dissections and scRNA-seq data analysis. L.L. and N.A.S. supervised the study. All authors discussed the results, contributed to data interpretation, revised the manuscript, and approved the final manuscript.

## Competing interests

The authors declare no competing interests.

## METHODS

### Mice

Mice were housed and bred in the animal facility at the Stowers Institute for Medical Research and handled in accordance with Institute and NIH guidelines. All procedures were approved by the Institutional Animal Care and Use Committee. C57BL/6J (CD45.2 mice), C57BL/6J-*Ptprc^em6Lutzy^*/J (CD45.1 mice), *Ncad-CreER*, VE-*cad-Cre*, *Cd200* ^fl/fl^, *Ncad-CreER; Cd200* ^fl/fl^, and VE-*cad-Cre; Cd200* ^fl/fl^ mice were used and maintained in a specific pathogen-free facility. All mouse strains were purchased from The Jackson Laboratory, except for the *Cd200* ^fl/fl^ mice, which were purchased from GemPharmatech Co., Ltd. (China). Animal rooms were maintained at 21-22 °C with 30-70% humidity under a 14 h light / 10 h dark cycle. Cages were changed every two weeks under a biological safety cabinet or laminar flow hood. For timed mating assays, mice were paired in the afternoon, and the morning on which a vaginal plug was detected was designated E0.5. Embryos were staged on the day of harvest by counting somite pairs.

### Flow cytometry and cell sorting

Flow cytometric analysis was performed using a BD ZE5 Cell Analyzer, and fluorescence-activated cell sorting was performed using a BD FACSymphony S6 cell sorter (BD Biosciences, NJ). Cells were stained with the indicated fluorophore-conjugated antibodies in 100 µL PBS containing 2% FBS. Where appropriate, dead cells were excluded by DAPI staining. Flow cytometry data were analyzed using FCS Express 7 software (De Novo Software, CA). The following antibodies were used at a dilution of 1: 200: VE-cadherin-BV421, CD45-PE/Cyanine5, TER119-BV421, CD41-PE/Cyanine5, CD31-PE/Cyanine7, c-KIT-APC-eFluor 780, CD144-PE, ESAM-FITC, CD45-APC, CD45.1-PE/Cyanine7, CD45.2-FITC, CD3E-PE, CD19-Alexa Fluor 700, CD11b (Mac-1)-PE/Cyanine5, Gr-1-APC, CD200-PE, N-cadherin-APC, CD200R1-PE, CD47-FITC, CD74-FITC, PD-L1-PE/Cyanine7, and CD73-PE-Cyanine7. Detailed information for each antibody is provided in Supplementary Table 2. Intracellular nitric oxide levels were measured by incubating cells in PBS containing 3 µM DAF-FM diacetate (Invitrogen, CA) for 20 min at 37 °C according to the manufacturer’s protocol. ERK phosphorylation was assessed using the ERK1/2 Phospho-PE/Cyanine7 (Thr202/Tyr204) antibody according to the manufacturer’s protocol. A complete list of primary and secondary antibodies is provided in Supplementary Table 2.

### Conditional knockout of *Cd200*

Conditional deletion of *Cd200* in N-cadherin-expressing cells was achieved by crossing Ncad-CreER; *Cd200* ^fl/fl^ males with *Cd200* ^fl/fl^ females. Pregnant females received an intraperitoneal injection of 2 mg tamoxifen at embryonic day (E) 9.5 to induce Cre-mediated recombination. Conditional deletion of *Cd200* in VE-cadherin-expressing cells was achieved by crossing VE-cad-Cre; *Cd200* ^fl/fl^ males with *Cd200* ^fl/fl^ females. Embryos were harvested at E10.5, E11.5, and E12.5 for analysis. Cre-mediated recombination was confirmed by RT-PCR analysis of embryonic brain tissue.

### AGM cell isolation

Mouse embryonic AGM tissues were dissected and enzymatically dissociated in 0.125% collagenase at 37 °C for 40 min, followed by mechanical dissociation. Cell suspensions were washed and filtered through a 40 μm cell strainer to obtain single-cell suspensions in PBS containing 2% FBS for subsequent analyses or transplantation assays. For flow cytometric analysis and cell sorting, freshly isolated AGM cells were stained with the indicated antibodies and analyzed or sorted using standard procedures.

### Transplantation assays

Six- to twelve-week-old female *Ptprc* recipient mice were lethally irradiated with a split dose of 9 Gy γ-irradiation (60Co) administered in two fractions separated by a 3 h interval. Donor cells were transplanted intravenously together with 5 × 10^5^ non-lysed CD45.1 splenocytes into irradiated *Ptprc* recipient mice. Peripheral blood was collected from the retro-orbital plexus using heparinized capillary tubes every 4 weeks until 16 weeks after transplantation to assess donor chimerism and multilineage distribution by flow cytometry. Recipients exhibiting ≥1% donor-derived chimerism were considered successfully engrafted.

### Immunofluorescence staining and imaging by confocal microscopes

E11.5 mouse embryos were fixed in 4% PFA at 4 °C overnight, followed by thorough washing with ice-cold PBS. For cryoprotection, embryos were incubated in 15% sucrose solution for 4 hours at room temperature, followed by incubation in 30% sucrose solution overnight at 4 °C. Embryos were then embedded in optimal cutting temperature (OCT) compound, and cryosectioned. Antigen retrieval was performed by heating the tissue sections in citrate buffer for 15 min at 70 °C. Tissue sections were then permeabilized and blocked before incubation with the indicated primary antibodies overnight at 4 °C. Sections were subsequently incubated with the appropriate fluorophore-conjugated secondary antibodies and counterstained with DAPI. After mounting, fluorescence images were acquired using a Nikon spinning disk microscope (Nikon Instruments, NY) equipped with a Plan Apo 20x/0.75 NA Air objective and a Hamamatsu ORCA-Fusion BT CMOS camera. Z-stack images were acquired with a pixel size of 0.325 µm, and a step size of 1 µm. Image processing and analysis were performed using Fiji and Imaris software (Bitplane, Switzerland).

### Whole-mount immunostaining and imaging

Embryos were prepared for whole-mount microscopy as previously described^52^. The CD31 primary antibody was used at a dilution of 1:500, and the c-KIT primary antibody was used at a dilution of 1:250. All secondary antibodies were used at a dilution of 1:1,000. Following immunostaining, embryos were optically cleared in benzyl alcohol/benzyl benzoate and imaged using a Leica confocal microscope (Leica Microsystems, Germany) (either an SP8 or Stellaris) equipped with a white light laser for excitation, a Plan Apo 20x/0.75 NA Oil objective, and HyD detectors for fluorescence collection. Z-stack images were acquired with a pixel size of 0.568 µm and a step size of 2 µm. Images were processed using Fiji, and three-dimensional reconstructions were generated in Fiji. CD31^+^ c-KIT^+^ IAHC cells were manually annotated in ImageJ and exported as ROI ZIP files. Using custom Python scripts, cell coordinates were projected onto an annotated 2D tube reference line, and distances were calculated relative to the vitelline artery within a 510-µm radius. To visualize cell density, relative distance values were grouped into contiguous spatial bins. The resulting binned counts were plotted to generate individual 2D spatial maps for each sample. To compare experimental groups, cell counts within corresponding distance bins were averaged across all samples in each group.

### Single-cell immunofluorescence analysis of β-catenin

Cells were fixed in 4% PFA for 30 min, washed three times with PBS, and applied to charged microscope slides by cytospin centrifugation (1,500 rpm, 3 min). Heat-induced antigen retrieval was performed in citrate buffer at 70 °C for 15 min. Cells were permeabilized with PBS-T containing Triton X-100, blocked, and incubated overnight at 4 °C with a mouse monoclonal anti-β-catenin antibody (Sigma-Aldrich, MO, C7207; 1:300). Cells were then incubated with goat anti-mouse Alexa Fluor 568 secondary antibody (Invitrogen, CA, A-11004) and counterstained with DAPI. Slides were mounted with ProLong Gold Antifade Mountant (Invitrogen, CA).

Images were acquired on a Leica STELLARIS confocal microscope (Leica Microsystems, Germany) using an HC PL APO CS2 63×/1.40 NA oil-immersion objective, with a pixel size of 0.05 µm and a z-step size of 0.3 µm. For quantification of the nuclear-to-cytoplasmic β-catenin fluorescence intensity ratio, individual cells were segmented using Cellpose, and nuclei were identified based on DAPI staining. The cytoplasmic region was defined as the area within each cell mask excluding the nuclear mask. Mean β-catenin fluorescence intensities were measured separately in the nuclear and cytoplasmic regions, and the nuclear-to-cytoplasmic ratio was calculated for each cell.

### Single-cell RNA sequencing

Fluorescence-activated cell sorting-enriched cells were assessed for concentration and viability using a Luna-FL cell counter (Logos Biosystems, South Korea). Samples with ≥80% cell viability were loaded onto a Chromium Single Cell Controller (10x Genomics, CA), and libraries were prepared using Chromium Next GEM Single Cell 3′ Reagent Kits v3.1, including either the Dual Index or Single Index workflow (10x Genomics), according to the manufacturer’s instructions. cDNA and short-fragment libraries were quantified and assessed for quality using a Qubit Fluorometer (Thermo Fisher Scientific, MA) and 2100 Bioanalyzer (Agilent Technologies, CA), respectively. Libraries representing ≥5,000 cells per sample were pooled and sequenced on Illumina NextSeq 500 or NovaSeq 6000 instruments (Illumina, CA), utilizing RTA and instrument software versions current at the time of processing, to achieve a mean sequencing depth of at least 40,000 reads per cell. Paired-end sequencing was performed using read lengths of 28×10×10×90 bp (dual index) or 28×8×91 bp (single index).

### Single-cell RNA sequencing data analysis

scRNA-seq data from seven samples (E10.5 AGM, E10.5 trunk, E11.5 AGM, E11.5 trunk, E12.5 AGM, E12.5 trunk, and E12.5 fetal liver) were processed using the Cell Ranger pipeline (version 6.0.1) with mouse mm10 Ensembl 98 annotation^53^, and the resulting filtered feature-barcode matrices were loaded into R (version 4.2.3) using the Seurat package (version 4.3.0)^54^. Gene names were converted to uppercase to ensure consistency across datasets. Cells were filtered based on the following quality control criteria: number of detected genes between 1,000 and 10,000, and mitochondrial gene content no greater than 5%. Filtered cells from all samples were merged into a single Seurat object. Gene expression counts were normalized using SCTransform with a Poisson regression model, fitting on up to 20,000 cells and selecting 3,000 variable features^55^. Principal component analysis (PCA) was performed using 100 components. UMAP was computed using the first 30 principal components for visualization. Shared nearest neighbor (SNN) graph-based clustering was performed using the first 30 principal components at multiple resolutions (0.2, 0.3, 0.5, and 0.8) to assess cluster stability. We used the clusters at a resolution of 0.8 for downstream analyses. Cluster-specific marker genes were identified using the receiver operating characteristic (ROC) test implemented in Seurat’s FindAllMarkers function, with a log fold-change threshold of 0.25, a minimum expression fraction of 10%, and retaining only positively enriched markers.

To contextualize our scRNA-seq data with known cell populations during embryonic hematopoiesis, we integrated our dataset with a previously published sorted AGM scRNA-seq dataset^36^. From the published dataset, the following cell populations were extracted: hemogenic endothelium (HE), venous endothelium (VE), arterial endothelium (AE), other endothelium, and intra-aortic hematopoietic clusters (IAHCs) from E10.5; pre-HSCs (E11.5 CD45^+^CD27^+^ IAHCs) and IAHCs from E11.5; and fetal liver HSCs (FL-HSC) from E14.5. Mitochondrial genes (prefixed with “MT^−^”) were removed from both datasets prior to integration. Gene expression matrices were subset to the intersection of genes detected in both datasets.

For integration, each dataset was independently normalized using log-normalization, and 3,000 highly variable features were identified per dataset using the variance-stabilizing transformation (VST) method. The union of variable features across datasets was used for downstream analysis. The merged dataset was log-normalized, scaled, and subjected to PCA (100 components). Batch correction was performed using Harmony (version 0.1)^56^, grouping by dataset of origin. UMAP embedding was computed using the first 50 Harmony-corrected dimensions. Integrated UMAP plots were generated to visualize the distribution of cell populations across datasets.

### Spatial transcriptomic profiling by Slide-seq V2 and Curio Seeker

Spatial transcriptomic profiling was performed using Slide-seq V2 or the Curio Seeker platform. For both approaches, mouse tissues were dissected, embedded in optimal cutting temperature compound to achieve transverse tissue sections and rapidly frozen at −90°C (HistoChill, Novec™ 7000; 3M, MN). Frozen tissues were cryosectioned at −12°C (Thermo, CryoStar NX70) at 10-µm thickness onto glass slides to locate region of interest prior to transferring sections directly onto the corresponding spatial transcriptomics capture substrate. Adjacent tissue sections were taken, fixed, stained and then imaged to use as references. The sections placed on the pucks/tiles did not undergo this process and were immediately placed in hybridization buffer for processing. For Slide-seq V2, spatially barcoded pucks and their associated barcode sequencing files were generated by the laboratory of Fei Chen at the Broad Institute according to the previously described Slide-seq V2 protocol^57^. Tissue sections were transferred directly onto the pucks, and mRNA capture, reverse transcription, tissue digestion, second-strand synthesis and cDNA amplification were performed according to the published protocol, incorporating minor modifications recommended by the Chen lab to optimize reverse transcription, bead handling and cDNA amplification. Sequencing libraries were generated from amplified cDNA using the Nextera XT library preparation workflow (Illumina, CA). Libraries were assessed for quality and concentration, pooled at equimolar ratios and sequenced on Illumina NextSeq platforms (Illumina, CA). Sequencing reads were demultiplexed to generate FASTQ files for downstream spatial transcriptomic analysis.

For experiments performed using the Curio Seeker platform, frozen tissue sections were transferred directly onto 3 × 3-mm Seeker tiles (Curio Biosciences, CA). Spatial RNA capture, cDNA synthesis and library preparation were performed according to the manufacturer’s protocol. Libraries were assessed for quality and concentration, pooled at equimolar ratios and sequenced on an Illumina NextSeq 2000 (Illumina, CA). Sequencing data were demultiplexed to generate FASTQ files for downstream analysis.

### Slide-seq V2 and Curio Seeker spatial transcriptomics data analysis

For each puck (Slide-seq V2) or tile (Curio Seeker), the gene expression count matrix and spatial bead coordinates were extracted, gene names were converted to uppercase, and barcodes were reformatted with stage and puck/tile identifiers. Beads with fewer than 50 total UMI counts were excluded. The filtered spatial data were normalized using SCTransform in Seurat (version 4.3.0) with a Poisson regression model, fitting on up to 20,000 beads and selecting 3,000 variable features. PCA was performed with 100 components.

Cell type identities were assigned to Slide-seq V2 / Curio Seeker beads using Robust Cell Type Decomposition (RCTD)^58^ implemented in the spacexr R package (version 2.2.1), which deconvolves the cell type composition of each spatial bead using a scRNA-seq reference. The scRNA-seq reference was prepared by subsetting the integrated scRNA-seq dataset to the matching developmental stage. For E10.5, endothelial populations (VE, AE, other endothelium) and IAC clusters from the published reference were excluded to avoid redundancy. For E11.5, IAC clusters from the published reference were similarly excluded. Cell type clusters with fewer than 25 cells were removed from the reference. The subsetted reference was re-processed with SCTransform (Poisson regression, 3,000 variable features) and PCA. RCTD was run in full deconvolution mode (doublet_mode = ‘full’), allowing mixed cell type assignments per bead, with a minimum UMI threshold of 50 per bead and a minimum of 25 cells per cell type in the reference. Cell type weights were normalized to sum to 1 for each bead.

To generate spatially resolved gene expression maps, scRNA-seq expression profiles were projected onto the Slide-seq V2 / Curio Seeker data. For each cell type cluster in the scRNA-seq reference, the mean log-normalized expression was computed across all cells. The projected expression for each bead was then calculated as a weighted linear combination of cluster-level mean expression profiles, using the RCTD-derived cell type proportions as weights (i.e., gene-by-cluster matrix multiplied by cluster-by-bead weight matrix).

To compare spatial transcriptomic patterns across developmental stages, one representative puck/tile from each stage (E10.5, E11.5, and E12.5) was integrated. Integration features (3,000 genes) were selected across SCTransform-normalized objects using SelectIntegrationFeatures, and scale data for these features were prepared with PrepSCTIntegration. After merging, PCA was performed on the SCT assay using 100 components, and batch effects across pucks were corrected using Harmony, grouping by puck identity. UMAP embedding was computed from the first 30 Harmony-corrected dimensions, and graph-based clustering was performed at multiple resolutions (0.2, 0.3, 0.5, and 0.8). We used the clusters at a resolution of 0.8 for downstream analyses. Spatial coordinates were adjusted with puck- or tile-specific rotation matrices for consistent anatomical orientation across stages. Cluster marker genes were identified using a logistic regression (LR) test with a log fold-change threshold of 0.1, a minimum expression fraction of 10%, and retaining only positively enriched markers.

### Cell-cell communication analysis on scRNA-seq data

Cell-cell communication was inferred from the scRNA-seq data using CellChat (version 2.1.2) with the CellChatDB.mouse ligand-receptor interaction database^59^. Analysis was performed independently at each developmental stage (E10.5, E11.5, and E12.5). For each stage, the scRNA-seq data were subset to AGM cells and their corresponding target populations: HECs at E10.5, pre-HSCs at E11.5, and E12.5 HSCs at E12.5 (cluster 14 cells along with fetal liver cells belonging to cluster 14). Gene expression was log-normalized, and gene symbols were mapped from uppercase back to their original case using an Ensembl 98 annotation table to match the CellChatDB nomenclature. Overexpressed ligands and receptors were identified per cell group, and overexpressed ligand-receptor interactions were computed. Gene expression was smoothed using the mouse protein-protein interaction (PPI) network. Communication probabilities were calculated using the trimean method on raw expression values, and interactions supported by fewer than 10 cells were filtered out. Pathway-level communication probabilities were computed, and interaction networks were aggregated.

To compare ligand-receptor interactions across developmental stages, communication probabilities for each source-target pair were extracted at a significance threshold of p < 0.05. Results were pivoted across stages, and for each ligand-receptor pair, the stage with the highest communication probability was identified. This enabled the characterization of temporally dynamic signaling patterns targeting HE, pre-HSCs, and their E12.5 counterparts. **Spatially aware cell-cell communication analysis.** To incorporate spatial proximity into cell-cell communication inference, CellChat (version 2.1.2) was applied to the Slide-seq V2 / Curio Seeker data using the projected gene expression profiles. Bead identities were assigned based on RCTD deconvolution predictions, with a minimum prediction score threshold of 0.25. Beads below this threshold were labeled as “Unassigned.“

Gene symbols were mapped to their original case using the Ensembl 98 annotation. The CellChat object was created in spatial mode, with bead coordinates provided directly. Spatial parameters were configured with a ratio of 0.73 and a tolerance of 5 (corresponding to a 10-micron bead diameter). Communication probabilities were computed using the trimean method with distance-dependent attenuation enabled (distance.use = TRUE), an interaction range of 250 microns, a distance scaling factor of 6.2, and contact-dependent interactions restricted to a range of 10 microns. Pathway-level communication probabilities and aggregated networks were computed as described above.

Interaction networks were visualized as both circle plots (showing interaction counts and strengths) and heatmaps using ComplexHeatmap (version 2.14.0)^60^. Spatially resolved communication networks were rendered by overlaying signaling edges on the Slide-seq V2 / Curio Seeker bead coordinates, with puck-specific rotation matrices applied for anatomical orientation. Node positions for each cell type were determined by the median spatial coordinates of all beads assigned to that type, with manual coordinate adjustments applied where needed for visual clarity.

### MERFISH processing and analysis

E11.5 mouse embryos were collected, embedded in optimal cutting temperature compound, and cryosectioned at 10 µm thickness. Tissue sections were processed for MERFISH using the MERSCOPE platform (Vizgen, MA) according to the manufacturer’s instructions, with minor modifications optimized for embryonic tissue. Briefly, sections were fixed, permeabilized, and hybridized with a custom MERFISH probe panel. Sections were imaged using the MERSCOPE system through sequential rounds of fluorescent readout hybridization, imaging, and signal removal. Raw imaging data were processed using the MERSCOPE analysis pipeline to decode individual RNA transcripts and assign them to their corresponding target genes, followed by cell segmentation and transcript assignment to individual cells. Processed spatial transcriptomic data were visualized and analyzed using MERSCOPE Visualizer and downstream computational analyses to characterize cell types, tissue organization, and regional gene-expression patterns within the E11.5 embryo.

### Data and code availability

Data and code will be available upon publication of the manuscript.

### Statistical analysis

All data are presented as means ± s.e.m. or means where applicable. Statistical analysis was performed by ANOVA, unpaired two-tailed t test or Fisher’s exact test unless otherwise indicated. Sequencing data were analyzed as described above. ns, not significant, P > 0.05; *P < 0.05; **P < 0.01; ***P < 0.001. The numbers of independent biological replicates are indicated in the figure legends.

---

**Extended Data Fig. 1.**
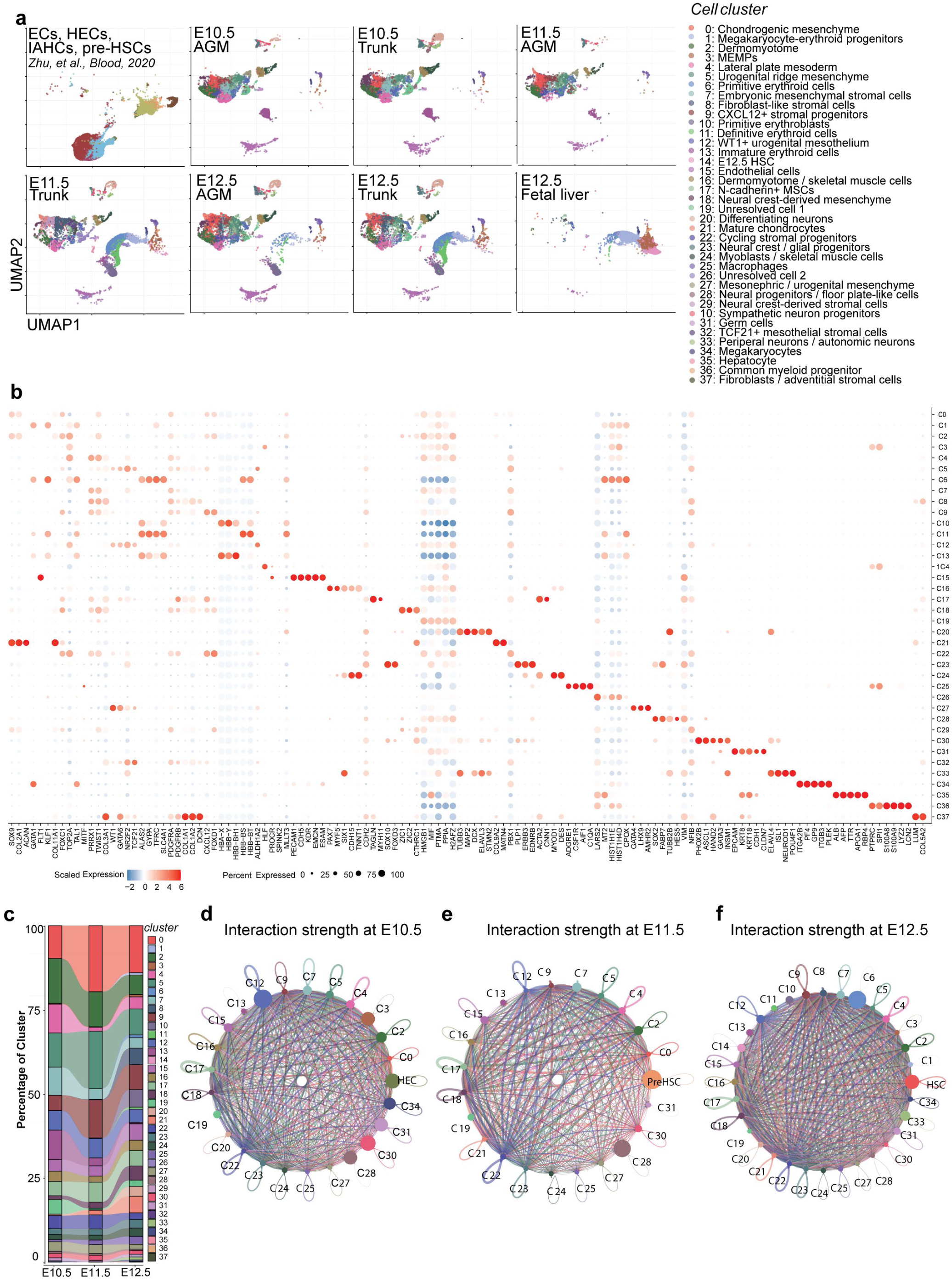
Single-cell transcriptomic atlas of the AGM. **a,** Integrated UMAP visualization of individual samples from the scRNA-seq datasets, comprising the published hematopoietic reference dataset and newly generated datasets from E10.5, E11.5 and E12.5 trunks; E10.5, E11.5 and E12.5 AGM; and E12.5 fetal liver. n = 3-5 biologically independent embryos for each group of newly generated datasets. **b,** Dot plot showing the expression of representative marker genes used for cell-type annotation. Dot size indicates the percentage of cells expressing each gene, and color represents the scaled gene expression. **c,** Relative abundance of annotated cell populations across E10.5, E11.5 and E12.5 AGM samples, illustrating dynamic changes in cell composition during HSC ontogeny. **d-f,** Cell-cell communication strength predicted using CellChat at E10.5 (**d**), E11.5 (**e**) and E12.5 (**f**). Circle size represents the relative number of inferred interactions associated with each cell population, and edge width indicates the predicted interaction strength between cell populations.

**Extended Data Fig. 2.**
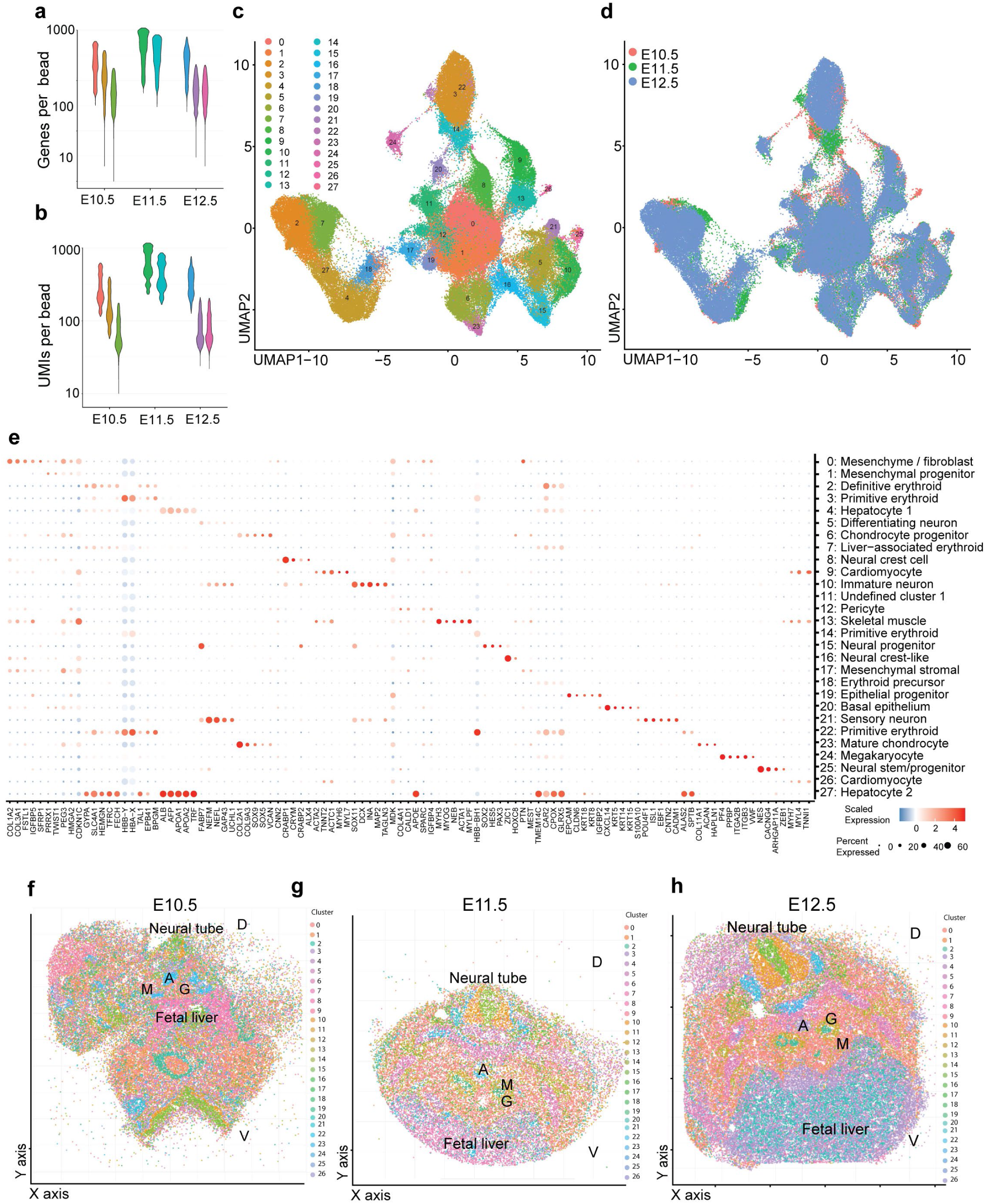
Quality assessment and clustering of Slide-seq V2 and Curio Seeker spatial transcriptomic data. **a, b,** Quality metrics of the Slide-seq V2 and Curio Seeker datasets across developmental stages. Violin plots show the number of detected genes per bead (**a**) and the number of unique molecular identifiers (UMIs) per bead (**b**) for E10.5, E11.5 and E12.5 samples. **c,** UMAP visualization of all Slide-seq V2 and Curio Seeker beads following unsupervised clustering, identifying 28 bead clusters. **d,** UMAP of the Slide-seq V2 and Curio Seeker beads colored by developmental stage (E10.5, E11.5 and E12.5), demonstrating successful integration of the datasets across stages. **e,** Dot plot showing the expression of representative marker genes used for annotation of the 28 Slide-seq V2 and Curio Seeker bead clusters. Dot size indicates the percentage of beads expressing each gene, and color represents the scaled gene expression. **f-h,** Spatial projection of annotated Slide-seq V2 or Curio Seeker bead clusters onto representative tissue sections from E10.5 (**f**), E11.5 (**g**) and E12.5 (**h**) embryonic trunks, reconstructing the anatomical organization of the embryonic trunk, including the AGM, neural tube and fetal liver. A, aorta; G, gonad; M, mesonephros; D, dorsal; V, ventral.

**Extended Data Fig. 3.**
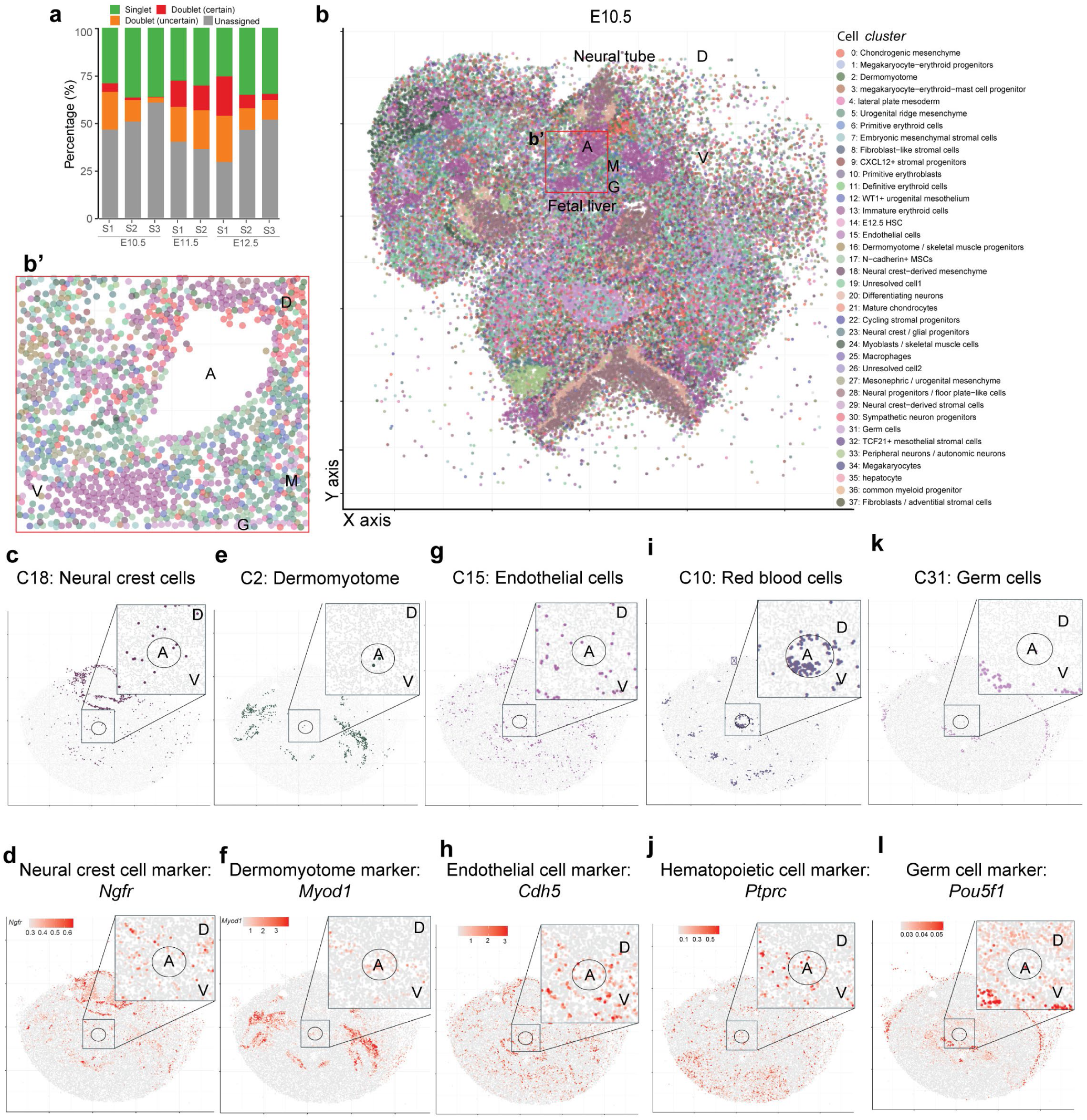
Integration of the scRNA-seq and the Slide-seq V2 and Curio Seeker datasets reconstructs the spatial organization of the niche in the AGM. **a,** Assignment of Slide-seq V2 and Curio Seeker beads following integration with the scRNA-seq reference dataset. Bars show the proportion of beads classified as singlets, uncertain doublets, certain doublets or unassigned for each biological replicate at E10.5, E11.5 and E12.5. **b,** Representative integrated spatial atlas of an E10.5 embryonic trunk. **b′,** Enlarged view of the boxed AGM region in **b**. For illustrative purposes, we removed erythroid cells in (**b′**) to better visualize the aortic structure. **c-l,** Representative spatial distribution of selected cell populations and their corresponding marker genes within the integrated spatial atlas at E11.5. **c, d,** Neural crest-derived mesenchyme and expression of the neural crest marker *Ngfr*. **e, f,** Dermomyotome and expression of the dermomyotome marker *Myod1*. **g, h,** endothelial cells and expression of the endothelial marker *Cdh5*. **i, j,** Erythroid cells and expression of the hematopoietic marker *Ptprc*. **k, l,** Germ cells and expression of the germ cell marker *Pou5f1*. A, aorta; G, gonad; M, mesonephros; D, dorsal; V, ventral. Circles represent the estimated position of the dorsal aorta.

**Extended Data Fig. 4.**
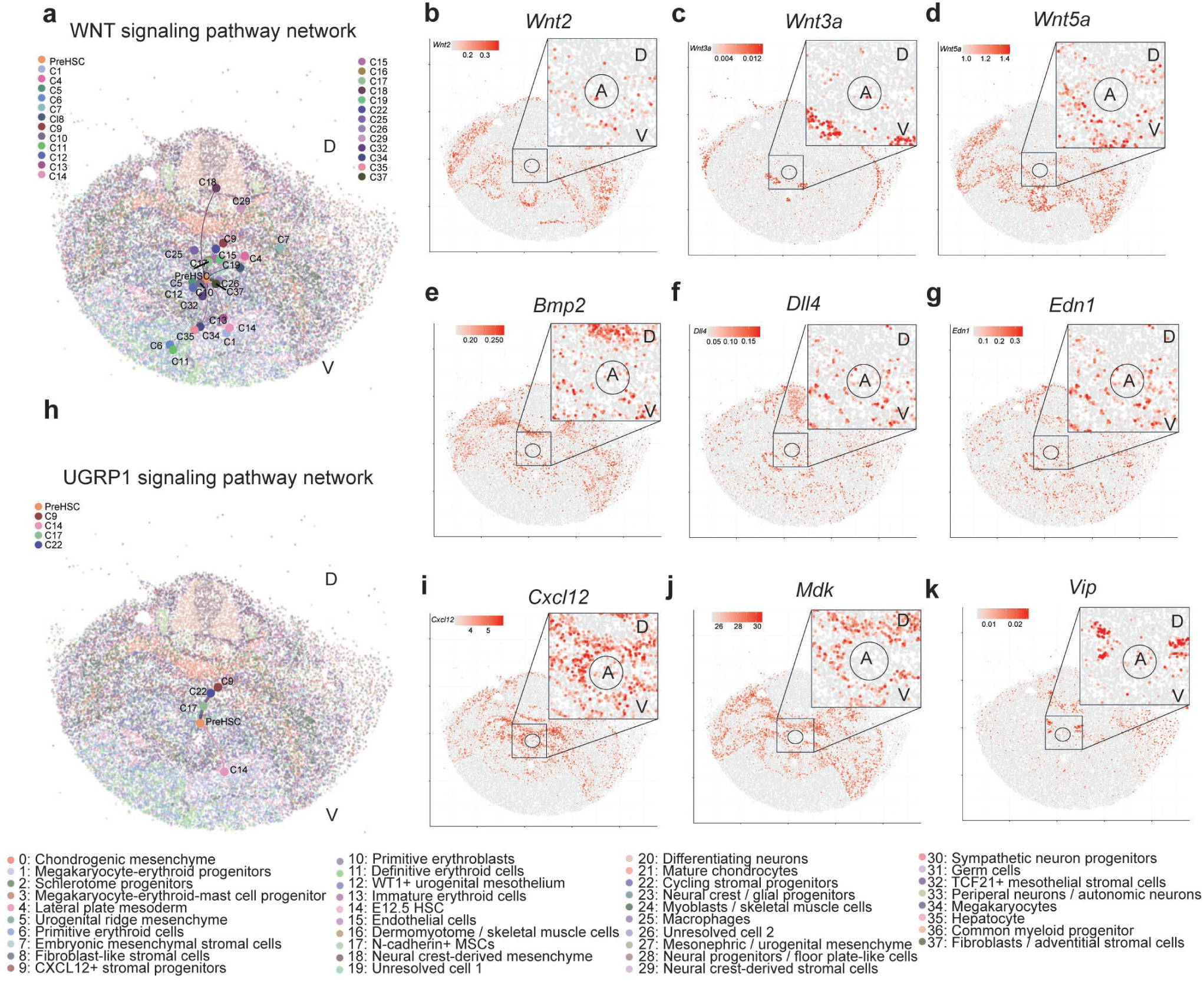
Spatial organization of additional developmental signaling pathways around the AGM. **a, h,** The integrated spatial atlas predicted spatially resolved interactions between cell populations and pre-HSCs via the WNT signaling network (**a**), and the UGRP1 signaling pathway network (**h**) within embryonic trunk sections at E11.5. Colored dots indicate ligand-producing cell populations. The position of each dot represents the center of the spatial distribution of the corresponding cell population in the Slide-seq V2 section. Line thickness indicates the interaction strength. **b-d,** Spatial distribution of representative WNT ligands within embryonic trunk sections at E11.5, including *Wnt2* (**b**), *Wnt3a* (**c**) and *Wnt5a* (**d**). **e-g, i-k,** Spatial distribution of representative signaling molecules within embryonic trunk sections at E11.5, including *Bmp2* (**e**), *Dll4* (**f**), *Edn1* (**g**), *Cxcl12* (**i**), *Mdk* (**j**), and *Vip* (**k**).

**Extended Data Fig. 5.**
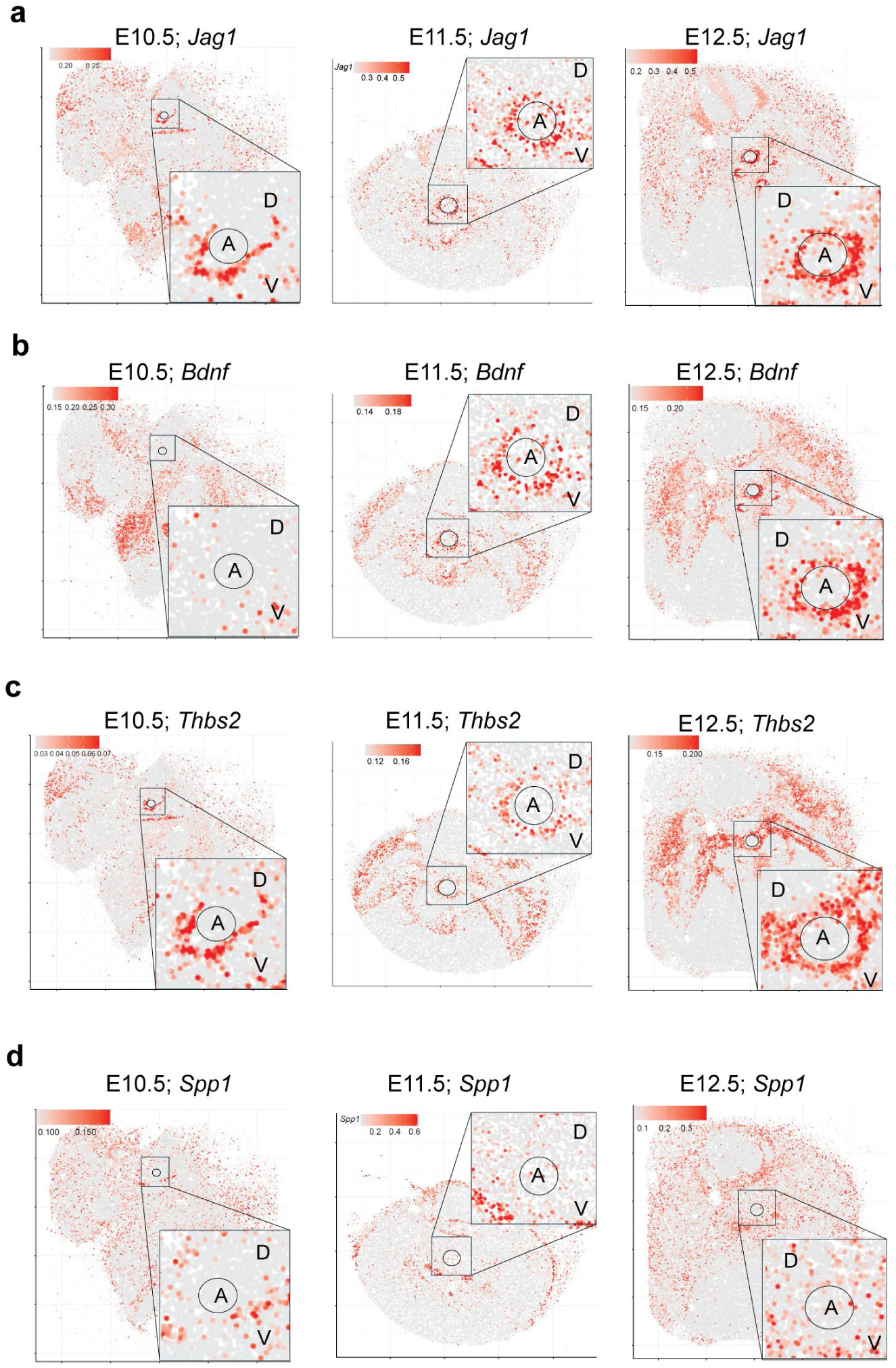
Dynamic spatial remodeling of niche-derived signaling during HSC ontogeny. **a-d,** Representative spatial expression of selected niche-derived signaling molecules in the integrated spatial atlas across developmental stages. Spatial expression of *Jag1* (**a**), *Bdnf* (**b**), *Thbs2* (**c**) and *Spp1* (**d**) within the embryonic trunk sections at E10.5, E11.5 and E12.5. Insets show enlarged views of the AGM region surrounding the dorsal aorta.

**Extended Data Fig. 6.**
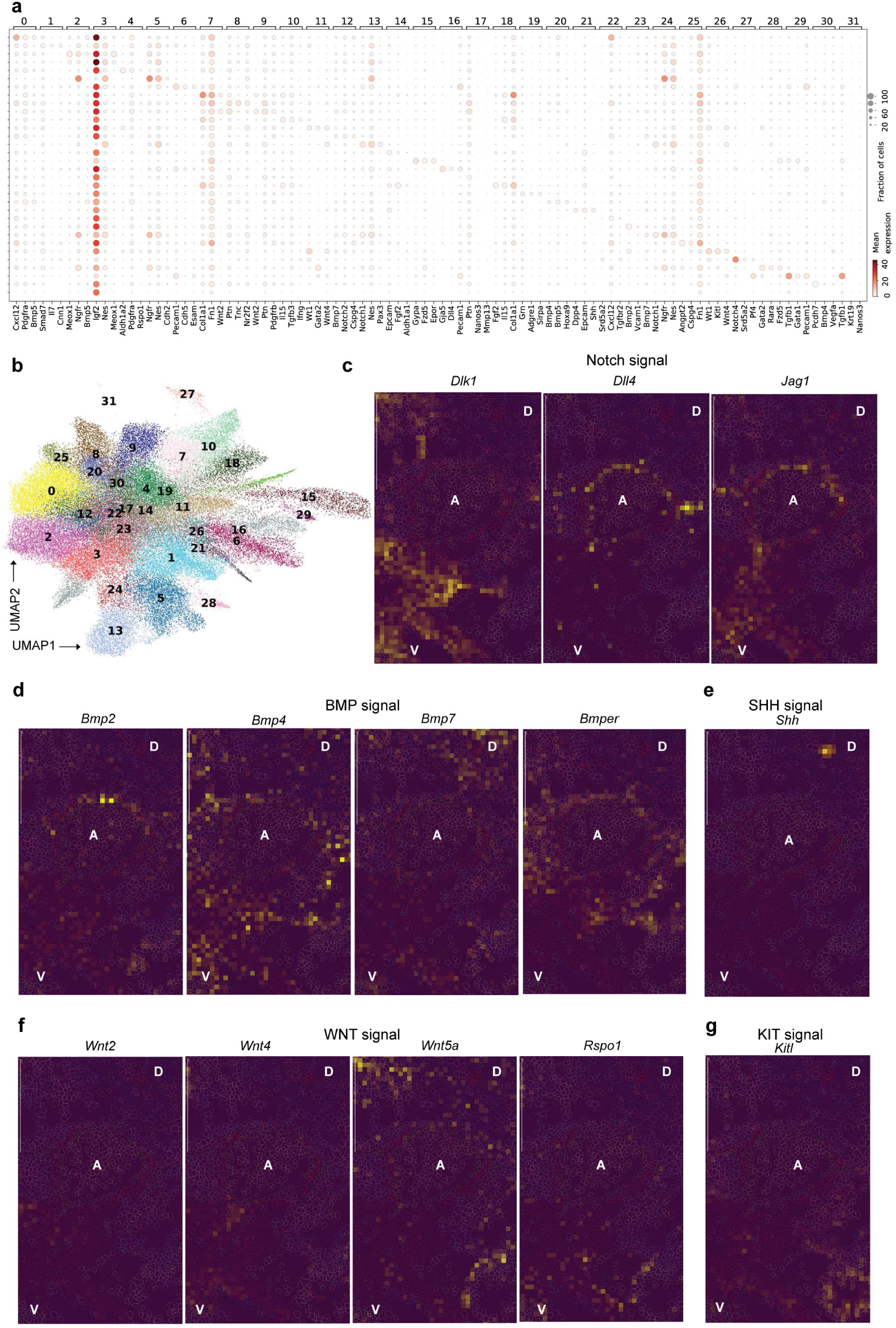
MERFISH validation of niche cell identity and spatial organization of developmental signaling. **a,** Dot plot showing the expression of representative marker genes across MERFISH-defined cell clusters. Dot size indicates the percentage of cells expressing each gene, and color represents the scaled gene expression. **b,** UMAP visualization of MERFISH-defined cell clusters. **c-g,** Representative MERFISH images showing the spatial expression of key developmental signaling molecules within embryonic trunk sections at E11.5. **c,** NOTCH signaling components (*Dlk1, Dll4* and *Jag1*). **d,** BMP signaling components (*Bmp2, Bmp4, Bmp7* and *Bmper*). **e,** *Shh* expression. **f,** WNT signaling components (*Wnt2, Wnt4, Wnt5a* and *Rspo1*). **g,** *Kitl* expression. A, aorta; D, dorsal; V, ventral.

**Extended Data Fig. 7.**
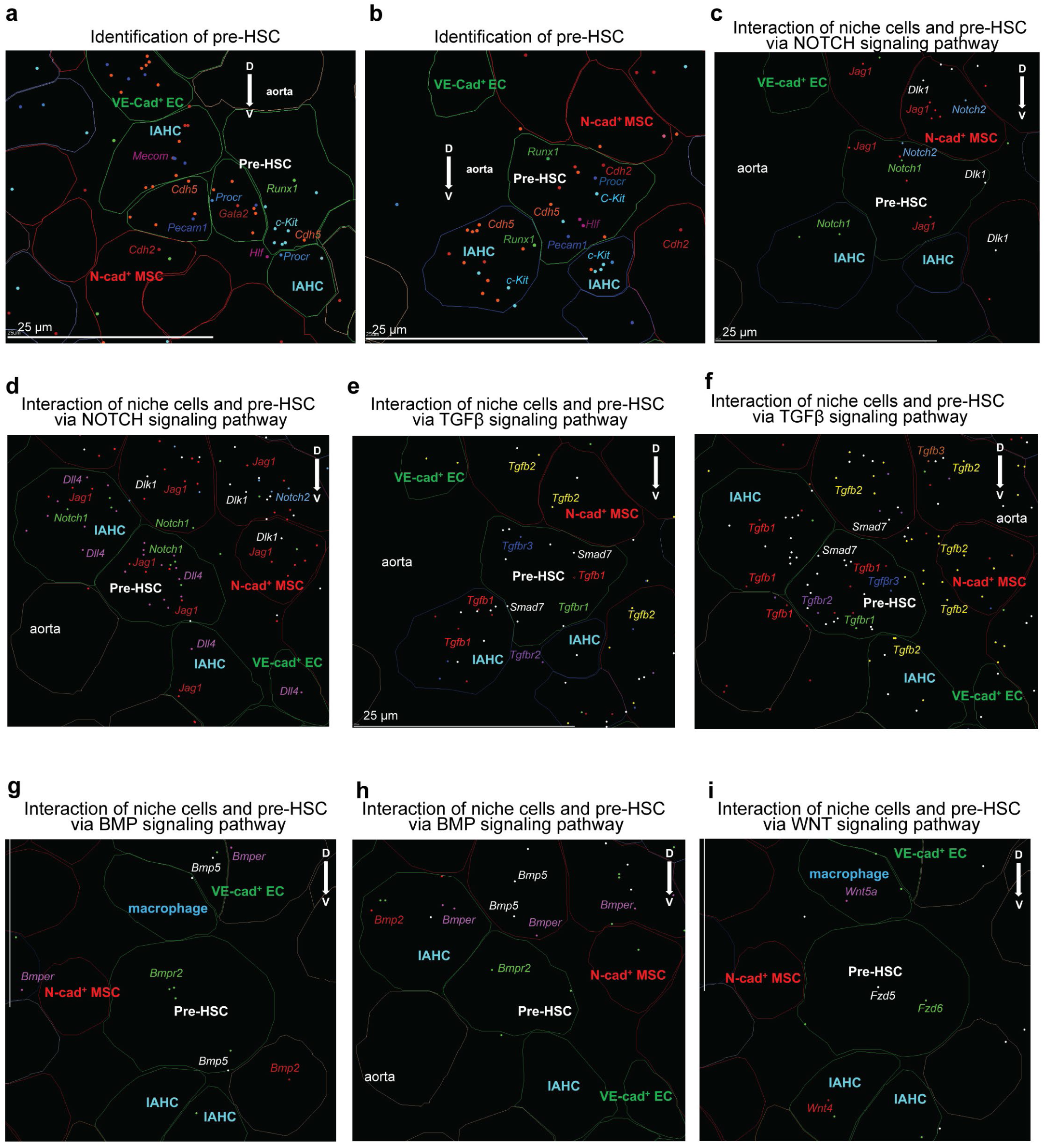
MERFISH identifies pre-HSCs in situ and visualizes signaling interactions within the native niche. **a, b,** Additional representative examples of individual pre-HSCs identified in situ by multiplex detection of pre-HSC-enriched transcripts. **c, d,** Additional representative MERFISH images showing ligand-receptor interactions between niche cells and pre-HSCs through the NOTCH signaling pathway. Ligands (*Jag1, Dll4* and *Dlk1*) and receptors (*Notch1* and *Notch2*) are shown in their endogenous spatial locations. **e, f,** Additional representative MERFISH images showing ligand-receptor interactions between niche cells and pre-HSCs through the TGFβ signaling pathway. Ligands (*Tgfb1-3*), receptors (*Tgfbr1-3*) and the downstream target *Smad7* are shown in their endogenous spatial locations. **g, h,** Representative MERFISH images showing the spatial organization of BMP signaling surrounding individual pre-HSCs. BMP ligands (*Bmp2, Bmp5* and *Bmper*) are detected in neighboring niche cells, whereas the receptor *Bmpr2* is expressed in pre-HSCs. **i,** Representative MERFISH image showing the spatial organization of WNT signaling surrounding a pre-HSC. The ligand *Wnt5a* is detected in neighboring niche cells, whereas the receptors *Fzd5* and *Fzd6* are expressed in the pre-HSC. A, aorta; D, dorsal; V, ventral. EC, endothelial cell. Each dot represents an individual transcript detected in situ. Scale bar = 25 µm.

**Extended Data Fig. 8.**
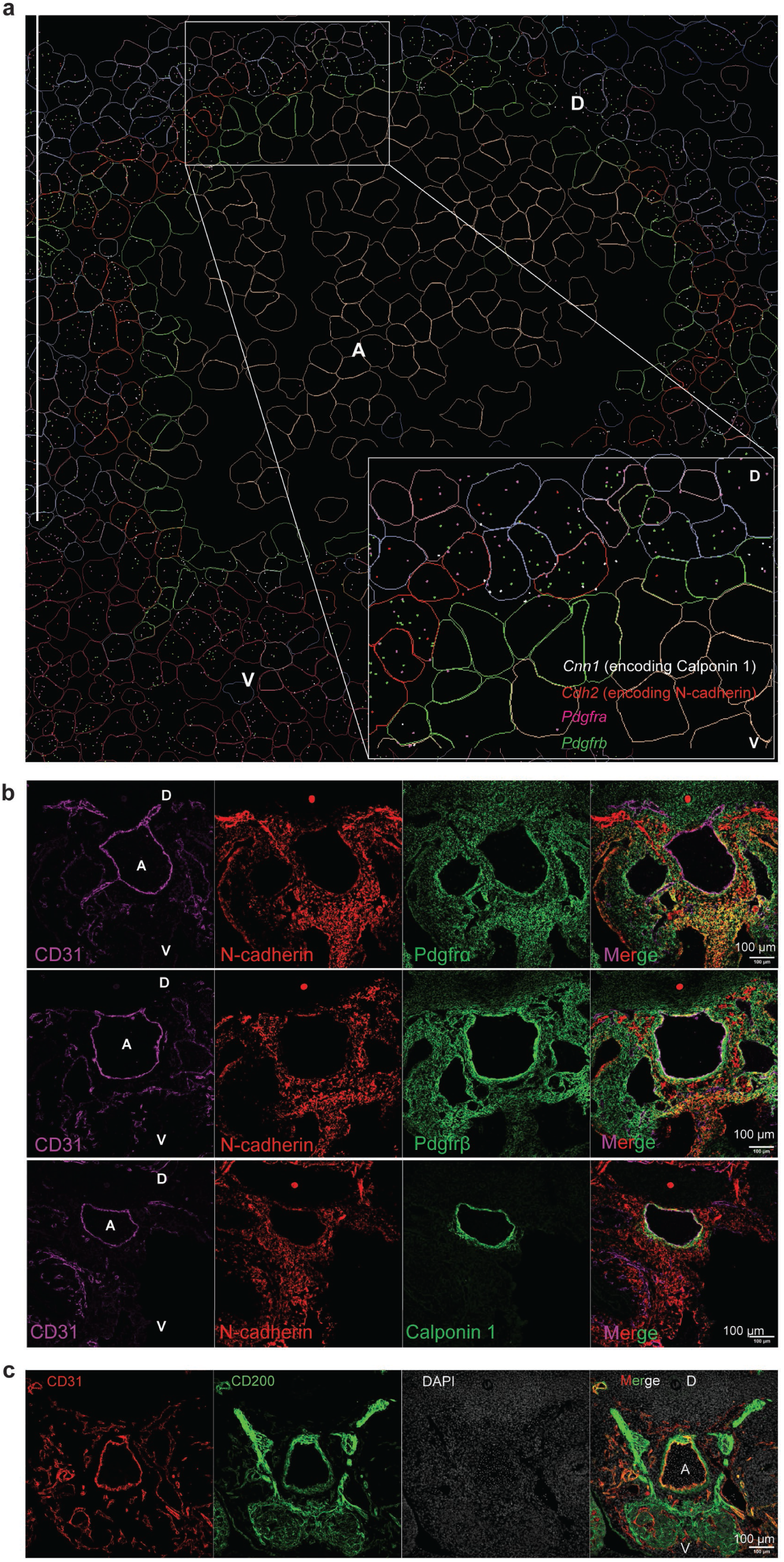
Characterization of N-cad⁺ MSCs and CD200 localization in the AGM. **a**, MERFISH validation of marker gene expression distinguishing N-cad⁺ MSCs from other peri-aortic mesenchymal populations. Shown are the spatial distributions of *Cdh2*, *Pdgfra, Pdgfrb* and *Cnn1*. **b**, Immunofluorescence validation of N-cad⁺ MSC identity. Representative images showing co-staining of CD31 (magenta), N-cadherin (red) and PDGFRα, PDGFRβ or Calponin-1 (green) on E11.5 embryonic trunk sections. **c**, Representative immunofluorescence staining of an E11.5 embryonic trunk section showing CD200 (green), CD31 (red) and DAPI (white). CD200 is enriched in the peri-aortic region surrounding the dorsal aorta and is associated with the aortic endothelial layer. n = 5 independent embryonic samples.A, aorta; D, dorsal; V, ventral. Scale bar = 100 µm in (**b, c**).

**Extended Data Fig. 9.**
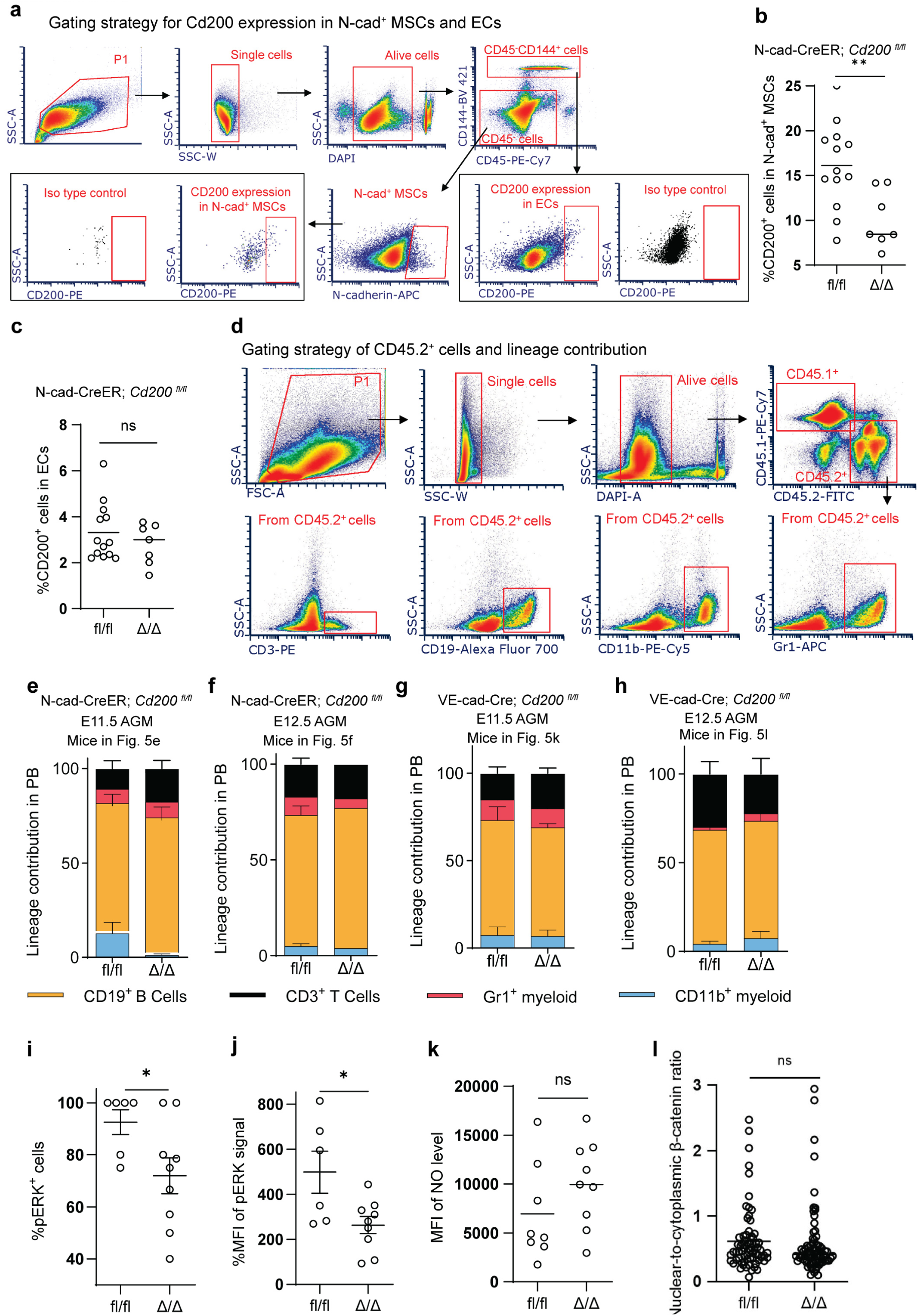
Validation of Cd200 conditional deletion and downstream functional analyses. **a,** Representative flow cytometry gating strategy used to quantify CD200 expression in N-cad⁺ MSCs and endothelial cells from E11.5 AGM. Live single cells were gated, followed by identification of CD45⁻CD144⁻ N-cad⁺ MSCs or CD45⁻CD144⁺ endothelial cells, and CD200 expression was quantified within each population. **b, c,** Quantification of the percentage of CD200⁺ cells among N-cad⁺ MSCs (**b**) and endothelial cells (**c**) from fl/fl and Δ/Δ embryos generated from *N-cad*-CreER; *Cd200* ^fl/fl^ mice. n = 13 for fl/fl group, and n = 7 for Δ/Δ group. **d,** Representative flow cytometry gating strategy for analysis of donor-derived hematopoietic reconstitution following transplantation. Live single donor-derived CD45.2⁺ cells were identified and further analyzed for lineage contribution using CD3 (T cells), CD19 (B cells), CD11b (myeloid cells) and Gr-1 (granulocytes). **e-h,** Lineage contribution of donor-derived peripheral blood cells following transplantation of AGM cells from the mice in Fig. 5e, f and Fig. 5k, l. Stacked bars show the relative contribution of CD19⁺ B cells, CD3⁺ T cells, Gr-1⁺ neutrophils and monocytes, and CD11b⁺ myeloid cells. **i, j,** Quantification of ERK phosphorylation in type II pre-HSCs isolated from fl/fl and Δ/Δ embryos at E12.5 from *N-cad*-CreER; *Cd200* ^fl/fl^ mice. **i,** Mean fluorescence intensity (MFI) of phosphorylated ERK (pERK). **j,** Percentage of pERK-positive cells. n = 6 for fl/fl group, and n = 9 for Δ/Δ group. **k,** Quantification of intracellular nitric oxide (NO) levels in type II pre-HSCs isolated from fl/fl and Δ/Δ embryos at E12.5 from *N-cad*-CreER; *Cd200* ^fl/fl^ mice, as measured by flow cytometry. n = 8 for fl/fl group, and n = 9 for Δ/Δ group. **l,** Quantification of the nuclear-to-cytoplasmic β-catenin ratio in type II pre-HSCs isolated from fl/fl and Δ/Δ embryos at E12.5 from *N-cad*-CreER; *Cd200* ^fl/fl^ mice, as assessed by immunofluorescence staining. Each dot represents one type II pre-HSC, from n = 4 for fl/fl group, and n = 4 for Δ/Δ group. Each dot in (**b, c, i-k**) represents one biologically independent embryo, and each dot in (**l**) represents one sorted type II pre-HSC. Lines indicate the means in (**b, c**) and means ± s.e.m. in (**e-l**). \**P* < 0.05; **P < 0.01; ns, not significant.

**Extended Data Fig. 10.**
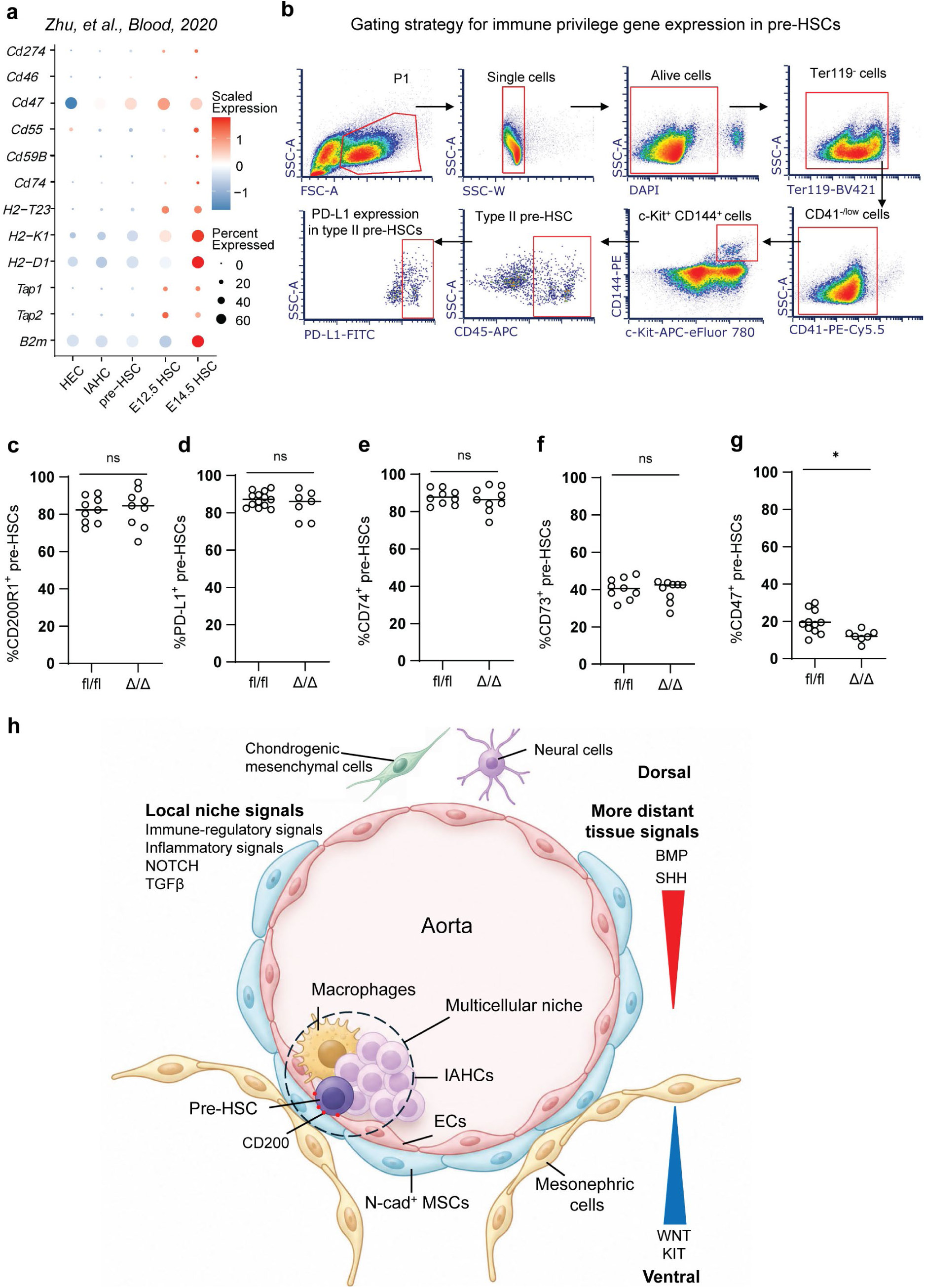
Progressive establishment of immune-regulatory characteristics during embryonic HSC maturation. **a,** Expression of representative immune-regulatory genes across hematopoietic developmental intermediates derived from the scRNA-seq dataset in Fig. 1a. Dot plot showing the scaled expression level (color) and percentage of expressing cells (dot size) for *Cd274* (encoding PD-L1), *Cd46, Cd47, Cd55, Cd59b, Cd74, H2-T23, H2-K1, H2-D1, Tap1, Tap2,* and *B2m* in HECs, IAHCs, pre-HSCs, E12.5 fetal liver HSCs, and E14.5 fetal liver HSCs. **b,** Representative flow cytometry gating strategy for quantification of immune-regulatory molecule expression in type II pre-HSCs at E12.5 AGM. Live single cells were sequentially gated after exclusion of Ter119⁺ erythroid cells. Type II pre-HSCs were identified as Ter119^−^CD41^−/low^ c-Kit⁺ CD144⁺ CD45⁺ cells. Representative gates for PD-L1 and CD47 expression are shown. **c-g,** Quantification of immune-regulatory molecule expression in type II pre-HSCs from littermate control (*Cd200* ^fl/fl^) and *N-cad-*CreER; *Cd200* ^Δ/Δ^ embryos. **c,** Percentage of CD200R1⁺ cells. **d,** Percentage of PD-L1⁺ cells. **e,** Percentage of CD74⁺ cells. **f,** Percentage of CD73⁺ cells. **g,** Percentage of CD47⁺ cells. Each dot represents one biologically independent embryo, and horizontal bars indicate the means. \**P* < 0.05; **P < 0.01; ns, not significant. **h,** Summary model of the spatial organization of the embryonic HSC niche in the AGM. Neural cells and chondrogenic mesenchymal cells are enriched dorsally, whereas N-cad⁺ MSCs, IAHCs, pre-HSCs, macrophages and mesonephric cells are enriched ventrally. BMP and SHH signaling show an overall dorsal enrichment, whereas WNT and KIT signaling are enriched ventrally. Pre-HSCs are embedded within a local multicellular niche comprising endothelial cells, N-cad⁺ MSCs, IAHCs and macrophages. Local niche signals include immune-regulatory, inflammatory, NOTCH and TGFβ signaling, whereas BMP, SHH, WNT and KIT signaling arise from more distant tissue compartments. CD200 is an EC-and N-cad⁺ MSC-derived contact-dependent immune-regulatory molecule that promotes HSC development.

## Notes

### Competing Interest Statement

The authors have declared no competing interest.

